# SRRM2 splicing factor modulates cell fate in early development

**DOI:** 10.1101/2023.12.15.571825

**Authors:** Silvia Carvalho, Luna Zea-Redondo, Tsz Ching Chloe Tang, Philipp Stachel-Braum, Duncan Miller, Paulo Caldas, Alexander Kukalev, Sebastian Diecke, Stefanie Grosswendt, Ana Rita Grosso, Ana Pombo

**Affiliations:** Max-Delbrück-Center for Molecular Medicine in the Helmholtz Association (MDC), Berlin Institute for Medical Systems Biology (BIMSB), Epigenetic Regulation and Chromatin Structure Group, 10115 Berlin, Germany; Associate Laboratory i4HB – Institute for Health and Bioeconomy, NOVA School of Science and Technology, Universidade NOVA de Lisboa, 2829-516 Caparica, Portugal; UCIBIO – Applied Molecular Biosciences Unit, Department of Life Sciences, NOVA School of Science and Technology, Universidade NOVA de Lisboa, 2829-516 Caparica, Portugal; Instituto de Ciências Biomédicas Abel Salazar (ICBAS), Universidade do Porto, 4050-313 Porto, Portugal; Graduate Program in Areas of Basic and Applied Biology (GABBA), ICBAS, University of Porto, 4050- 313 Porto, Portugal; Humboldt-Universität zu Berlin, Institute of Biology, Berlin, Germany; Berlin Institute of Health (BIH) at Charité – Universitätsmedizin Berlin, 10178 Berlin, Germany; Max-Delbrück-Center for Molecular Medicine in the Helmholtz Association (MDC), Berlin Institute for Medical Systems Biology (BIMSB), From Cell State to Function Group, 10115 Berlin, Germany; Max-Delbrück-Center for Molecular Medicine in the Helmholtz Association (MDC), Pluripotent Stem Cells Platform, 13125 Berlin, Germany; DZHK (German Centre for Cardiovascular Research), partner site Berlin, 10785 Berlin, Germany

**Author notes:** Shared co-corresponding Lead contacts and correspondence: Prof. Ana Pombo, Max-Delbrück-Center for Molecular Medicine in the Helmholtz Association (MDC), Berlin Institute for Medical Systems Biology (BIMSB), Hannoversche Str. 28, 10115 Berlin, Germany. Prof. Ana Rita Grosso, NOVA School of Science and Technology, Campus de Caparica, 2829-516 Caparica, Portugal. Co-authors (Luna Zea-Redondo, Tsz Ching Chloe Tang).

**Keywords:** Srrm2, SRm300, splicing, pluripotency, stemness, single-cell transcriptomics

## Abstract

Embryo development is an orchestrated process that relies on tight regulation of gene expression to guide cell differentiation and fate decisions. Alternative splicing is modulated during development as an additional layer of regulation to reprogram gene expression patterns. The *Srrm2* splicing factor has recently been implicated in developmental disorders and diseases, but its role in early mammalian development remains unexplored. Here, we show that *Srrm2* dosage is critical for maintaining embryonic stem cell pluripotency and cell identity. *Srrm2* heterozygosity promotes loss of stemness, characterized by the coexistence of cells expressing naive and formative pluripotency markers, together with extensive changes in gene expression, including genes regulated by serum-response transcription factor and differentiation-related genes. Depletion of *Srrm2* by RNA interference in embryonic stem cells shows that the earliest effects of Srrm2 half-dosage are specific alternative splicing events on a small number of genes, followed by expression changes in metabolism and differentiation-related genes. Our findings unveil molecular and cellular roles of *Srrm2* in stemness and lineage commitment, shedding light on the roles of splicing regulators in early embryogenesis, developmental diseases and tumorigenesis.

**Summary statement:** This article emphasizes the importance of splicing regulators in early mammalian development by uncovering roles of SRRM2 splicing factor dosage in pluripotency, providing novel insights for a better understanding of Srrm2-related diseases.

## Introduction

Embryo development is a dynamic complex process that requires refined spatial and temporal gene expression regulation, coordinating cell differentiation and cell fate decisions. A major aspect of gene expression that contributes to gene function diversity is alternative splicing. Changes in the expression levels of specific splicing factors can induce cell differentiation of pluripotent cells and commitment to specific embryonic lineages. Specific splicing events of developmental transcription factors can promote and modulate differentiation (Tsai et al., 2014; Mayshar et al., 2008; Venables et al., 2013; Gabut et al., 2011; Han et al., 2013). Genome-wide programs of alternative splicing are also dynamic during cell differentiation or cell lineage reprogramming (Baralle and Giudice, 2017; Wu et al., 2010; Revil et al., 2010; Cloonan et al., 2008). Although alternative splicing is increasingly understood as a dynamic process that accompanies gene expression rewiring during development, the specific roles of splicing regulators remain ill defined.

Splicing is catalysed by small nuclear ribonucleoproteins (snRNPs) and non-small nuclear ribonucleoproteins splicing factors. The pan-eukaryotic serine/arginine repetitive matrix (SRRM) family of SR-related proteins are splicing factors that lack RNA-binding domains and play important roles in both constitutive and alternative splicing regulation (Blencowe et al., 1998; Torres-Méndez et al., 2019). The SRRM family comprises four elements, SRRM1, SRRM2, SRRM3 and SRRM4. Both SRRM1 and SRRM2 promote early stages of splicing, by mediating interactions between SR proteins bound at exonic enhancers and at splice or branch sites marked by U1/U2 snRNPs, typically promoting exon inclusion (Blencowe et al., 1998; Eldridge et al., 1999; Blencowe et al., 1999).

SRRM1 and SRRM2 proteins are broadly expressed among different tissues, while SRRM3 and SRRM4 are expressed in brain tissues. Misregulation of SRRM2 has been implicated in cancer and neurological disorders (Tomsic et al., 2015; Shehadeh et al., 2010; Cuinat et al., 2022). SRRM3/4 play crucial roles in microexon splicing, with important roles in brain function (Irimia et al., 2014; Quesnel-Vallières et al., 2016), and their misregulation can lead to neurological and neurodevelopmental problems (Nakano et al., 2019; Calarco et al., 2009; Quesnel-Vallières et al., 2016; Nakano et al., 2012; Quesnel-Vallières et al., 2015). Strikingly, a recent meta-analysis study of 31,058 human exomes identified SRRM2 as a novel gene linked with developmental disorders, displaying the strongest enrichment for *de novo* mutations (Kaplanis et al., 2020). *SRRM2* expression decreases during the differentiation of human primary macrophages and its depletion upregulates differentiation marks in human myeloid-like cells (Liu et al., 2018; Xu et al., 2022).

*SRRM2* has a high probability of intolerance to loss-of-function variants in humans (Karczewski et al., 2020; Petrovski et al., 2013), suggesting that it plays a gene-dosage dependent critical role in key biological processes. Data from the International Mouse Phenotyping Consortium also identify *Srrm2*-dosage effects in *Srrm2*-knockout mice, whereby homozygous loss-of-function leads to embryonic lethality at the pre-weaning stage, whereas *Srrm2*-heterozygous mice are viable (www.mousephenotype.org, Groza et al., 2022). *Srrm2* roles in embryonic development have been reported for its *C. elegans* ortholog (Fontrodona et al., 2013), but not yet investigated in mammalian development.

In this study, we use constitutive and transient depletion of *Srrm2* in mouse embryonic stem cells (mESC) as a tool to gain biological insights into *Srrm2* roles in early mammalian development. We found that *Srrm2* dosage is critical for maintaining embryonic stem cell pluripotency and that its downregulation alters pluripotent states and cell identity. We show that half dosage of *Srrm2* leads to specific splicing alterations that precede extensive changes in gene expression and perturbation of the stemness phenotype in mESC.

## Results

### *Srrm2* reduction impairs the maintenance of mESC pluripotency

To explore the expression of *Srrm1/4* genes in early development, we mined single-cell and bulk gene expression data from public resources (https://apps.kaessmannlab.org/evodevoapp/, https://endoderm-explorer.com/, Cardoso-Moreira et al., 2019; Nowotschin et al., 2019). We found that *Srrm2* is highly expressed in E3.5-5.5 blastocyst cells, with slightly increased expression from E3.5 inner cell mass (ICM) cells to E5.5 visceral endoderm (VE) cells and especially epiblast cells (**Fig. 1A**). *Srrm2* is also expressed in all tissue lineages, albeit with different expression levels and with a tendency to decrease, especially in non-neuronal tissues, during embryonic development (**Fig. 1B**).

**Fig. 1.**
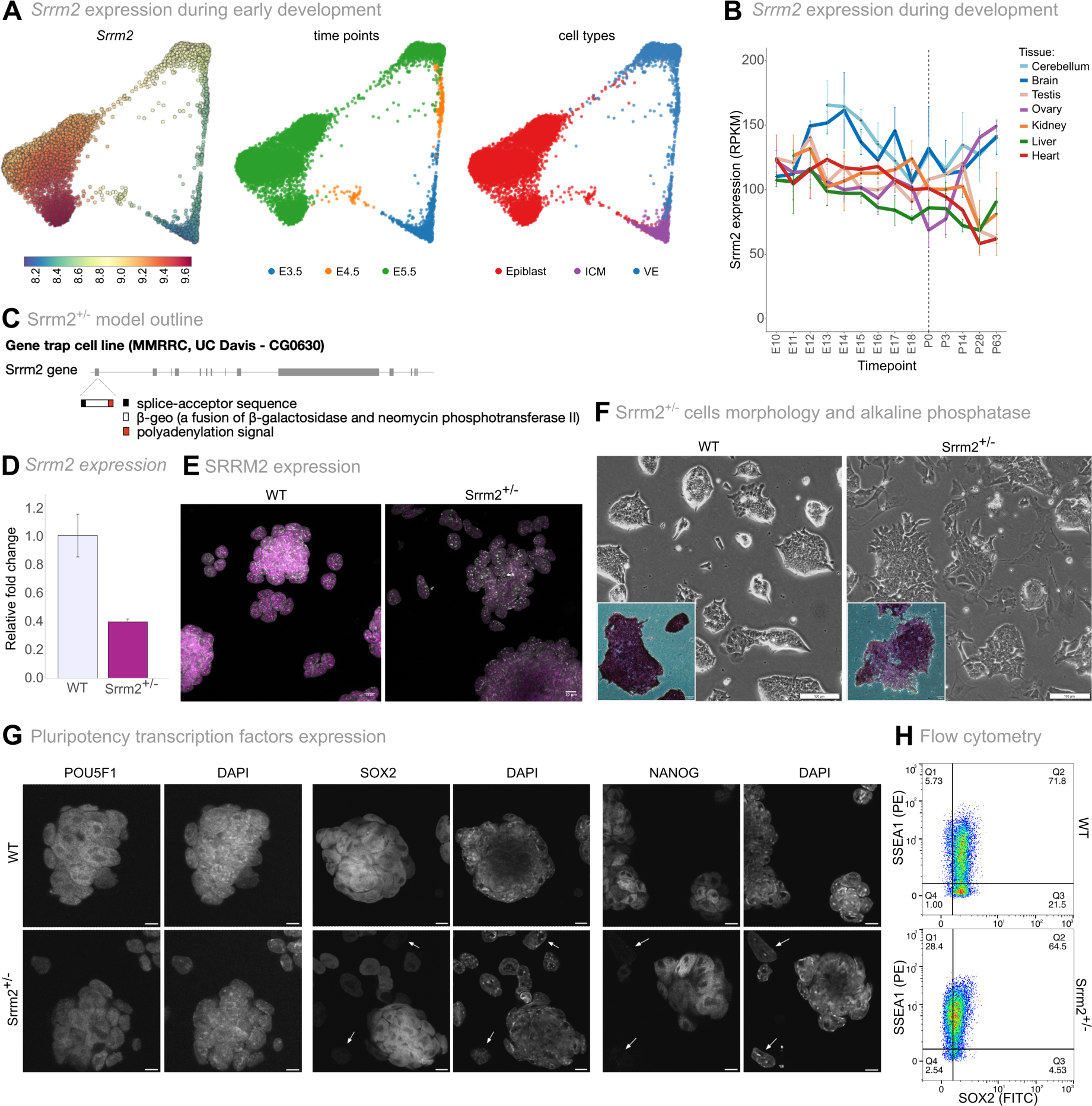
*Srrm2* expression levels influence the pluripotent state of mESC. A) Single-cell expression of *Srrm2* during E3.5-E5.5 mice embryo development (Nowotschin et al., 2019). Force- directed layouts coloured by *Srrm2* gene expression (left), development time (middle) or cell type labels (right; ICM: inner cell mass; VE: visceral endoderm). **B)** *Srrm2* expression levels during pre- natal (E) and post-natal (P) developing mouse tissues (RPKM) (Cardoso-Moreira et al., 2019). The vertical dashed line represents the birth time point. **C)** Srrm2^+/-^ knockout gene-trap embryonic stem cell line description. Gene-trap cassette was inserted in Exon 1 of the *Srrm2* gene. **D)** *Srrm2* expression levels. Total RNA levels of exon 11 of *Srrm2* were measured by RT-qPCR. Relative levels are normalized to β-actin. Mean and standard deviation are calculated from three biological replicates. **E)** Immunofluorescence of SRRM2 in WT and Srrm2^+/-^ cell cultures. Depicted by a representative experiment of at least 3 independent experiments from 3 biological replicates. Each image corresponds to the merge of the DAPI (grey) and SRRM2 (purple) channels. Scale bars correspond to 10 µm. **F)** Brightfield images from WT and Srrm2^+/-^ cell cultures and alkaline phosphatase staining of WT and Srrm2^+/-^ cells. Brightfield images: representative experiment data of at least 3 independent experiments from 3 biological replicates. Scale bar corresponds to 100 µm. Insets: Alkaline phosphatase staining of WT and Srrm2^+/-^ cells. Representative experiment data from 3 biological replicates in at least 3 independent experiments. Scale bar corresponds to 10 µm. **G)** Immunofluorescence of POU5F1, SOX2 and NANOG pluripotency transcription factors (left to right) in WT and Srrm2^+/-^ cell cultures (above to below). Each image corresponds to an individual channel per immunofluorescence (first: pluripotent transcription factor - POU5F1, SOX2 or NANOG; second: DAPI). Representative images from two independent immunofluorescence experiments for SOX2 and from one experiment for POU5F1 and NANOG, performed in one biological replicate. Scale bars correspond to 10 µm. Arrows indicate cells with very little or absent expression of the respective pluripotency factor. **H)** Flow cytometry data of SSEA1 and SOX2 double labelling in WT and Srrm2^+/-^ cells. Data shown is representative of 1 out of 3 biological replicates.

To investigate the effects of *Srrm2* dosage in pluripotency and cell lineage commitment, we obtained a gene-trap mouse stem-cell line with a heterozygous truncation in exon 1 of the *Srrm2* gene (Srrm2^+/-^) from the Mutant Mouse Resource Center (MMRRC), originally generated by the Sanger Institute Gene Trap Resource (SIGTR) (**Fig. 1C**). *Srrm2* transcript levels measured by qRT-PCR showed a 60% reduction in Srrm2^+/-^ cells compared to the parental wildtype ESC line (WT; **Fig. 1D**). Immunofluorescence experiments showed that SRRM2 protein levels are homogeneously reduced in the Srrm2^+/-^ cell population **(Fig. 1E)**.

To test whether *Srrm2* depletion affects the ESC pluripotency phenotype, we started by assessing the morphology of cell colonies using brightfield imaging of WT and Srrm2^+/-^ cells grown in a serum-free medium supplemented with leukemia inhibitory factor (LIF). We found that Srrm2^+/-^ cells formed fewer well-defined colonies than WT cells, grew in dispersed clusters of cells, and showed an increased number of cells outside colonies (**Fig. 1F**). We also measured alkaline phosphatase (AP) activity, and found that the majority of Srrm2^+/-^ cells were positive for AP activity, but showed lower signal intensity than WT cells (**Fig. 1F insets**). The cell clusters and cells outside colonies were typically low or negative in AP activity in Srrm2^+/-^ cells, suggesting a change in the regulation of the pluripotent state (Štefková et al., 2015).

To further investigate the state of pluripotency of Srrm2^+/-^ cells, we performed immunofluorescence of classic pluripotency-associated factors POU5F1, SOX2 and NANOG, and found that they were similarly expressed in WT and Srrm2^+/-^ cells within the colonies or clusters (**Fig. 1G**). Consistent with a change in the pluripotent state, the isolated cells outside colonies or cell clusters, expressed lower levels of the pluripotency-associated factors, and were observed more frequently in Srrm2*^+/-^*cell cultures (**Fig. 1G**, arrowheads point to cells with undetectable expression). Western blot analyses confirmed that NANOG is expressed in Srrm2^+/-^ cells, although in lower bulk levels (**Fig. S1A**). We found that AP intensity and the expression of pluripotency markers remained constant in the three biological replicates of Srrm2^+/-^ cell cultures with increased passage. The visual appearance of Srrm2^+/-^ cell cultures did not noticeably change throughout the study, suggesting that the partial loss of colony integrity and altered heterogenous profile is stable.

To better understand the balance between cell states within the Srrm2^+/-^ and WT cell cultures, we performed flow cytometry of SOX2 and of the pluripotency-associated cell surface marker SSEA1 (Cui et al., 2004). We found that Srrm2^+/-^ cells generally express lower levels of SOX2 than WT cells, with an increase in the number of SSEA1-positive cells (**Fig. 1H**, **Fig. S1B**). These results show that the loss of colony formation in Srrm2^+/-^ cells is not trivially related to loss of pluripotency, but may represent an intermediate pluripotent stage. Moreover, the lower dosage of *Srrm2* is sufficient to induce a stable culture state characterized by fewer ESC that express pluripotency markers, which nevertheless retain the ability to form colonies.

### *Srrm2* heterozygous knockout leads to extensive gene expression changes

To further characterize the Srrm2^+/-^ cells, we performed total RNA profiling using high-throughput sequencing (RNA-seq) in three biological replicates. We found extensive changes in gene expression between Srrm2*^+/-^* and WT cells, with 2569 upregulated and 2205 downregulated genes (adj. P-value < 0.05; **Fig. 2A, Table S1**). We confirmed that *Srrm2* was downregulated in Srrm2^+/-^ cells (fold change of 1.61, adj. P-value 2.36e-05, **Fig. S2A**, **Table S1**). The expression of *Nanog* and *Klf4,* the first core pluripotency factors to be downregulated as cells exit pluripotency, were reduced by 1.7 and 1.8 fold, respectively, whereas *Sox2* was reduced by 1.2 fold, and *Pou5f1* was not differentially expressed (**Fig. S2B, Table S1**). These results are consistent with a partial loss of stemness of the Srrm2*^+/-^* cells, as hinted by the initial imaging characterization, with similarities to intermediate states of pluripotency described in mouse and human cells (Kalkan and Smith, 2014; Smith, 2017; Messmer et al., 2019). Amongst the top differential expressed genes, we found many associated with embryo development (e.g., *Mest*, *Itga3* and *H19),* including gametogenesis (e.g., *Ddx3y* and *Eif2s3y).* Most differentially expressed genes were protein coding, with a small tendency for upregulation (**Fig. S2C**). We also found downregulation of 103 out of 232 snoRNAs (**Fig. S2D**), including *Gm12238*, *Snora65* and *Snora24,* implicating *Srrm2* in the expression or processing of snoRNAs.

**Fig. 2.**
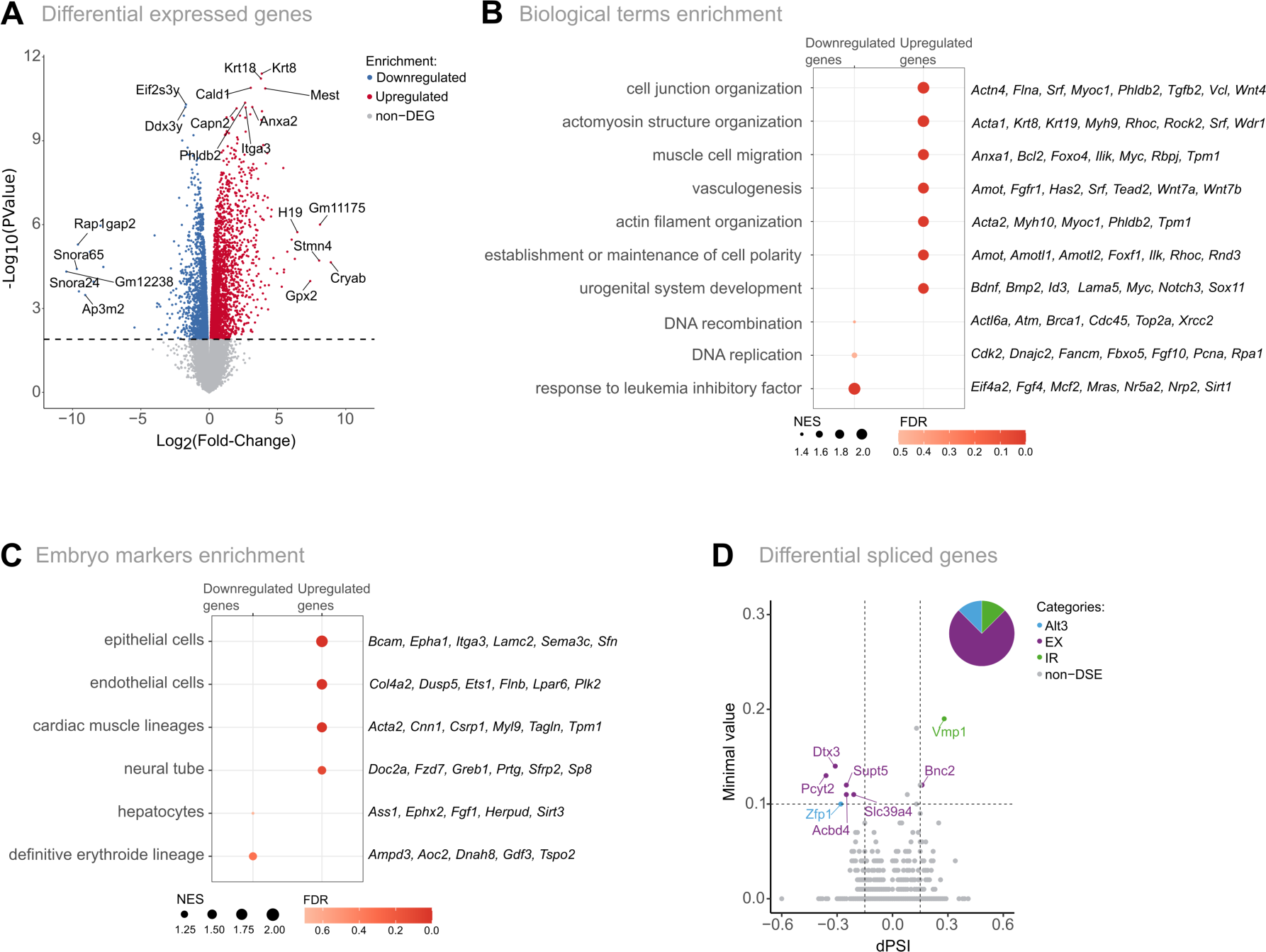
*Srrm2* heterozygous knockout mESC show extensive gene expression alterations. A) Volcano plot showing the Srrm2^+/-^ differentially expressed genes in total RNA from 3 replicates of each condition. 18,912 genes were detected in the whole dataset and 4,774 genes were considered as differentially expressed, using an adjusted P-value < 0.05 as a cut-off (downregulated and upregulated genes are marked in blue and red, respectively). Gene labels represent the top-10 differentially expressed genes with the lowest adjusted P-value, and the top-5 differentially expressed genes with the higher absolute fold change. **B)** GSEA for Gene Ontology (GO) biological processes of Srrm2^+/-^ relative to WT gene expression comparison. Red intensity and the size of the circles represent FDR values and Normalized Enrichment Score (NES), respectively. **C**) GSEA for publicly available E9.5-13.5 single-cell embryo tissue markers (Cao et al., 2019) for comparison of Srrm2^+/-^ versus WT gene expression. Red intensity represents the FDR values, whereas the circle size indicates the Normalized Enrichment Score (NES), respectively. **D**) Volcano plot showing the differentially spliced events between Srrm2^+/-^ and WT cells, where the magnitude of alterations is represented by the difference in Percent Spliced In (dPSI) and the minimal value, computed by Vast- tools. 261,655 splicing events were detected in the whole dataset and 8 splicing events were considered as significantly differentially spliced, using a cut-off of abs(dPSI) ≥ 0.1 and a minimal value ≥ 0.1. Spliced events with a dPSI < 0 correspond to exclusion events in Srrm2^+/-^ cells, whereas dPSI > 0 correspond to inclusion events in Srrm2^+/-^ cells. Categories of significant differential spliced events are represented by different colours: alternative 3’ splice site (Alt3, blue), alternative 5’ splice site (Atl5, orange), exon exclusion (EX, purple), intron retention (IR, green), no alteration (non-DSE, grey). The pie chart represents the proportion of significant differential spliced events.

Gene set enrichment analysis (GSEA) focused on biological processes, showed that up-regulated genes in Srrm2^+/-^ cells are associated with cytoskeletal-related terms and differentiation processes, namely mesoderm development, as vasculogenesis (e.g., *Srf*, *Amot*, *Wnt7a*, *Wnt7b*) and urogenital development (e.g., *Sox11*, *Podxl*, *Ca2*, *Plaur*; **Fig. 2B, Table S2**). In contrast, down-regulated genes were significantly associated with response to LIF (e.g., *Nr5a2*, *Nrp2*, *Mras*, *Mcf2*), which plays a central role in promoting a pluripotent ground state. Together with the imaging results, the bulk transcriptomics data show that Srrm2^+/-^ cells have a mixed expression signature with features of pluripotency and differentiation.

### Srrm2 heterozygous knockout leads to the acquisition of diverse differentiation signatures

To further explore whether the differentiation signature of Srrm2^+/-^ cells has a preferred developmental lineage, we performed a GSEA using publicly available single-cell transcriptome profiles from mouse embryos (Cao et al., 2019). We found that Srrm2^+/-^ upregulated genes were significantly enriched for markers of epithelial, endothelial and cardiac muscle cells (FDR adj. P-value < 0.05; **Fig. 2C, Table S3**), while down-regulated genes included markers of hepatocyte and definitive erythroid lineages (adj. P-value > 0.05).

We noticed that many differentiation transcription factors were up-regulated in Srrm2^+/-^ cells, including *Srf*, *Mest* and *Sox11* (**Fig. 2A-B**). Over-representation analysis (ORA) using transcription factor lists with the potential to drive transdifferentiation, as predicted by Mogrify (Rackham et al., 2016; Ferrai et al., 2017), showed that Srrm2^+/-^ upregulated genes are preferential enriched for transcription factors associated with differentiation into cardiomyocytes, fibroblasts and mesenchymal cells (**Fig. S2E**, **Table S4**). Transcription factor network analysis showed that many upregulated genes are targets of the Serum Response Factor (SRF) (**Fig. S2F-G, Table S5**), which is one of the top 15 genes encoding for transcription factors with the lowest P-value (**Fig. S2H**). SRF is involved in cell differentiation, controls cell-to-cell adhesion, expression of cytoskeleton genes during development, and is required for epithelial and cardiac differentiation (Verdoni et al., 2010; Posern and Treisman, 2006; Miano et al., 2007; Deshpande et al., 2022). These results indicate that Srrm2^+/-^ cells have a transcriptional profile consistent with partial loss of pluripotency combined with the expression of differentiation markers that represent different cellular states or lineage specialization.

### Srrm2 heterozygous knockout shows specific splicing alterations

Given the known roles of SRRM proteins in splicing regulation, we next asked whether the extensive gene deregulation of Srrm2^+/-^ cells was generally associated with splicing defects. We detected differential spliced events in the total RNA-seq data using Vast-tools (Han et al., 2017; Irimia et al., 2014; Braunschweig et al., 2014; Tapial et al., 2017), and identified only eight alternatively splicing events and affected genes, with the majority corresponding to exon exclusion events (**Fig. 2D, Table S6**). Six out of the eight genes were also differentially expressed (*Bnc2, Dtx3, Slc39a4* were downregulated, while *Pcyt2*, *Vmp1* and *Zfp1* were upregulated), and five play important roles in embryo development. VMP1 is important for autophagy and cell adhesion and is involved in lipid metabolism and in embryo implantation (Ropolo et al., 2007; Guo et al., 2012; Morishita et al., 2019; Holzner et al., 2023). DTX3 is a ubiquitin ligase that regulates NOTCH signaling, with key roles in development (Ding et al., 2020; Wang et al., 2021b). BNC2 and PCYT2 are important for the differentiation and maturation of mesenchyme and muscle, respectively, both with origin in the mesodermal germ layer (Vanhoutteghem et al., 2009; Zhu et al., 2009). *Pcyt2* and *Slc39a4* are essential genes for mouse development, as complete knockout mice die during embryogenesis (Fullerton et al., 2007; Pastor-Arroyo et al., 2021). Our results reveal that the transcriptional profile of Srrm2^+/-^ cells is consistent with an altered pluripotent state that affects several development-related genes and is compatible with a mixture of different cellular states within the Srrm2^+/-^ cell cultures.

### Srrm2^+/-^ cell populations display heterogeneity of pluripotency states

To further explore the transcriptional and pluripotent state of Srrm2^+/-^ cells, we generated single-cell RNA-seq (scRNA-seq) data using 10x Genomics technology from two biological replicates of WT and Srrm2^+/-^ cell cultures. Following quality control processing, we obtained a total of 31,433 cells and a median of 4,097 genes detected per cell. The WT and Srrm2^+/-^ single-cell transcriptome data was integrated using the Harmony algorithm, revealing four distinct cell clusters, represented through dimensionality reduction via uniform manifold approximation and projection (UMAP; **Fig. 3A**). Most WT cells (86%) are grouped in cluster 1, whereas 94% of Srrm2*^+/-^* cells populate clusters 2 to 4 (**Fig. 3B**; **Fig. S3A**), suggesting that *Srrm2* heterozygosity gives rise to a distinct cell state. In line with the immunofluorescence analyses of SRRM2 protein expression, we found that *Srrm2* was expressed at lower levels in Srrm2^+/-^ cells independently of the state (**Fig. S3B**). Cycling cells were also found in all clusters, with clusters 1 and 2 having similar proportions of G1/G0, S and G2 cells, while cluster 4 shows the largest proportion of G1/G0 cells (**Fig. S3C)**. These analyses confirm that Srrm2^+/-^ cells have a distinct and heterogeneous transcriptional profile which reflects the morphological variability observed in the cell cultures.

**Fig. 3.**
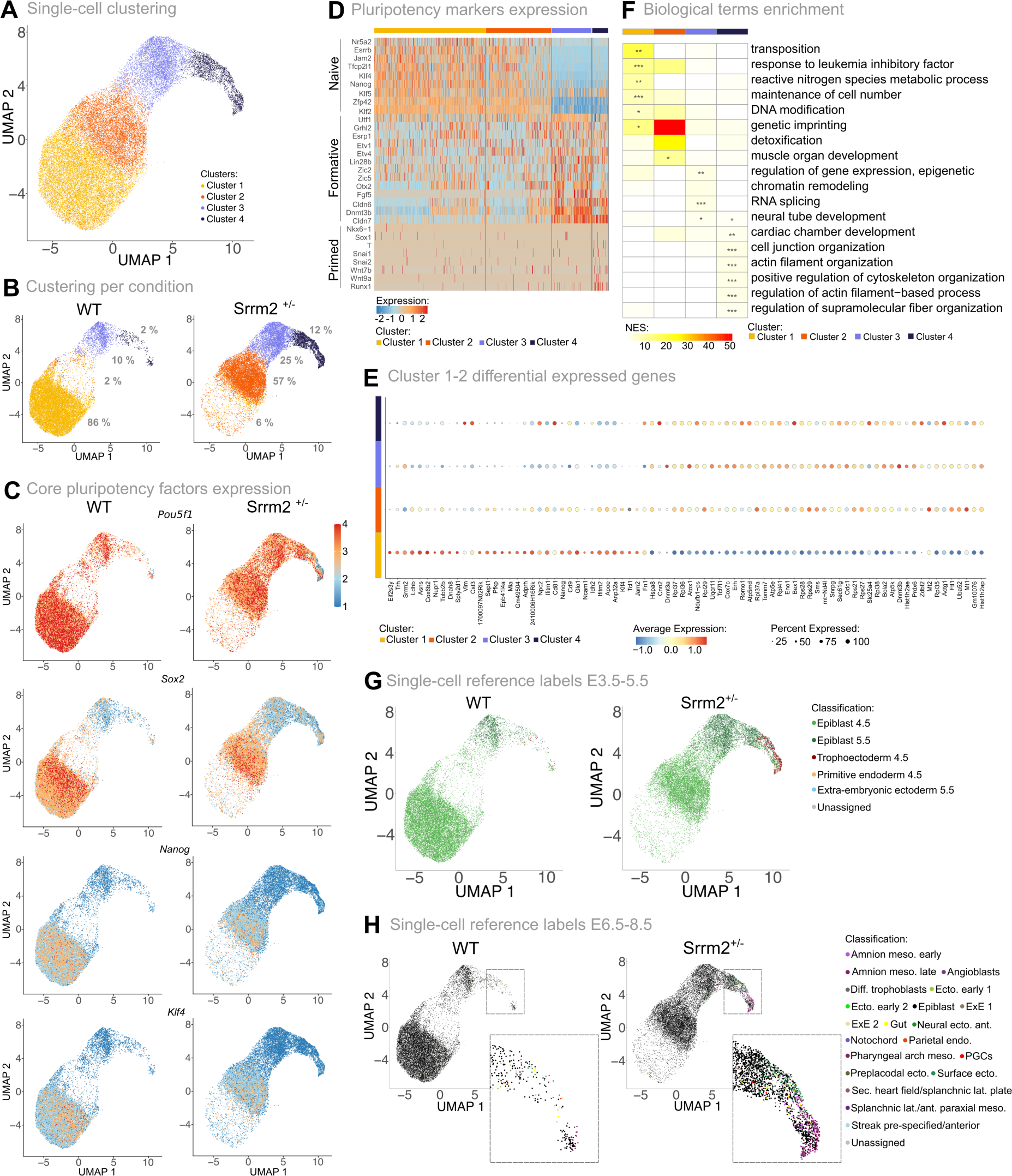
Single-cell transcriptome alterations in *Srrm2* heterozygous knockout mESC. **A-C)** Uniform Manifold Approximation and Projection (UMAP) graphs represent the relative similarity between individual cells: coloured by cell cluster (**A**), shown separately for WT or Srrm2^+/-^ cells (**B**), or coloured by the expression levels of core pluripotency factors (**C**). The displayed numbers in (B) represent the percentage of cells in each cluster per condition. **D)** Expression of a panel of pluripotency markers for naive, formative and primed states along the different cell clusters. Gene markers are labelled on the left side. Each line represents the expression in a single cell. Cells are divided into clusters (from left to right, clusters 1 to 4), illustrated by different colours on the top of the heatmap. **E)** Dotplot representing the expression levels of differentially expressed genes between cells of clusters 1 and 2 per cluster. Clusters are represented in different colours on the left of the plot (from down to top, clusters 1 to 4). Circle size corresponds to the percentage of cells expressing the gene of interest. Colour scale represents the average level of expression per cluster. **F)** Heatmap representing biological GO terms enrichment from Over- Representation Analysis (ORAs) using gene markers per cluster. The top 5 significant terms per cluster are displayed. Normalized enrichment scores (NES) of GO terms are represented by a continuous colour scale with yellow representing lowest enrichment and red highest enrichment. Significant NES values are marked with an asterisk (Fisher exact test *: adjusted P-value < 0.05; **: adjusted P-value < 0.01; ***: adjusted P-value < 0.001). **G-H)** UMAP projection coloured according to the predicted transferred ID labelling from publicly available *in vivo* scRNA-seq reference atlas of mouse embryogenesis cells from E3.5-5.5 (Nowotschin et al., 2019) (G) and E6.5-8.5 (Chan et al., 2019; Grosswendt et al., 2020, Haggerty et al., 2021) (H). Cells are classified into amnion mesoderm early, amnion mesoderm late, angioblasts, differentiated trophoblasts, ectoderm early 1, ectoderm early 2, epiblast, extraembryonic ectoderm 1 (ExE 1), extraembryonic ectoderm 2 (ExE 2), gut, neural ectoderm anterior, notochord, parietal endoderm, pharyngeal arch mesoderm, primordial germ cells (PGCs), preplacodal ectoderm, surface ectoderm, secondary heart field/splanchnic lateral plate, splanchnic-lateral/anterior-paraxial mesoderm, streak pre-specified/anterior, unassigned (H).

We next assessed the expression of pluripotency transcription factors in the different cell states and confirmed that *Pou5f1*, *Sox2*, *Nanog* and *Klf4* are less expressed in Srrm2^+/-^ than in WT cells (**Fig. 3C**). *Pou5f1* is expressed in all clusters, although in lower levels in Srrm2^+/-^ cells from cluster 4 (**Fig. S3D**). *Sox2* is also expressed in all clusters, but at lower levels in clusters 3 and 4, while *Nanog* and *Klf4* expression is not detected in most cells in clusters 3 and 4 (**Fig. S3D**). These data show that clusters 1 and 2 resemble the most stem-like states in WT and Srrm2^+/-^ cultures, whereas clusters 3 and 4, which are more abundant in Srrm2*^+/-^* cell cultures, are characterized by a lower expression of pluripotency transcription factors.

Given the heterogeneous expression profiles of the core pluripotent markers, we asked whether Srrm2^+/-^ cells depict previously characterized pluripotency states, namely naive (capable of giving rise to all embryo lineages), formative (intermediate state initiating specialization to a particular lineage), or primed (induction of specialized markers) (Kalkan and Smith, 2014; Smith, 2017; Messmer et al., 2019). We measured the expression of markers of the different pluripotency states (Wang et al., 2021) in the four different clusters and found that naive state markers are expressed in clusters 1 and 2, although at higher levels in cluster 1 (**Fig. 3D**). Formative markers are most highly expressed in clusters 3 and 4, whereas primed state markers are mostly not detected, except for *Wnt9a* and *Runx1* in cluster 4. Together, these data reveal that Srrm2^+/-^ cell cultures contain cells in different states of pluripotency, including an intermediate pluripotent state that displays shared features between the naive and formative pluripotency, and other states that are characterized by the absence of pluripotency signatures.

The exit of pluripotent stem cells from the pluripotent naive state is accompanied by alterations in gene expression networks, increased DNA methylation, and metabolism changes, resulting in cell fate determination (Hoogland and Marks, 2021; Kim and Costello, 2017). To further investigate the intermediate pluripotent state of Srrm2*^+/-^* cells, we quantified the transcriptional differences between clusters 1 and 2, and detected 35 downregulated genes and 45 upregulated genes (**Fig. 3E, Table S7**), from which 50 genes were also differentially expressed in the bulk data, but not differentially spliced. Amongst the downregulated genes, we found *Srrm2*, *Nanog* and *Klf4*, but also the mitochondrial protein *Cox6b2* with previously described roles in the stemness of trophoblast cells (Saha et al., 2022). Within the upregulated genes, we found the transcription factor *Tcf7l1*, which mediates the transition from naive to primed states of pluripotency by repressing the expression of naive and formative genes, and drives cells towards primitive endoderm through the Wnt/β-catenin signalling pathway (Guo et al., 2011; Hoffman et al., 2013; Pereira et al., 2006; Athanasouli et al., 2023). Other up-regulated genes included *de novo* DNA methyltransferases, *Dnmt3a* and *Dnmt3b*, which have roles in maintaining the naive pluripotency state (Li et al., 2017; Kinoshita et al., 2021; Betto et al., 2021; Leitch et al., 2013), and ribosomal protein genes, in line with previous reports of global translation increase in early differentiation and cell fate transitions (Li and Wang, 2020).

Finally, to further characterize the different clusters, we identified cluster-specific gene markers (**Table S8**) and performed Gene Ontology (GO) enrichment analyses (**Table S9**). The group of gene markers of the WT-linked cluster 1 was enriched for genes associated with response to LIF (e.g., *Klf4*, *Zfp42*) and stem cell pluripotency (e.g., *Nanog*, *Sox2, Esrrb1, Rest, Lefty2*) (**Fig. 3F**). The most stem-like group of Srrm2^+/-^ cells (cluster 2) has 23 gene markers, which include *Dppa3*, *Zfp42*, but not *Klf4*, *Nanog* or *Sox2*, as well as ribosomal subunits and genes associated with muscle organ development (e.g., *Rbpj, Tcf15*, *Mymx*). Finally, cluster 3 gene markers were enriched for terms such as epigenetic remodellers, neural tube and cardiac chamber development (e.g., *Dnmt3a*, *Dnmt3b*, *Arid1a*, *Sox4* and *Sox11*), whereas cluster 4 was mostly enriched for cytoskeleton-related terms.

Interestingly, cluster 3 and 4 markers are associated with embryonic lethality and abnormal development, including cardiovascular and blood vessel morphology (**Fig. S3E, Table S10**). Finally, we also observed that the expression of SRF target genes was mostly enriched in cluster 4, despite no detectable changes in *Srf* gene expression (**Fig. S3F**). These results suggest that Srrm2*^+/-^* cells have combined features associated with partial loss of pluripotency into more advanced stages of early development and specific embryonic lineages.

To investigate a potential equivalence between Srrm2^+/-^ cells and *in vivo* embryonic development, we leveraged published scRNA-seq data of mouse embryogenesis covering either the pre- implantation blastocyst into an implanted egg cylinder stage (E3.5-5.5; Nowotschin et al., 2019), or the gastrulation and early organogenesis stages (E6.5-8.5; Chan et al., 2019; Grosswendt et al., 2020). We transferred cell-type annotations from the two references, E3.5-5.5 and E6.5-8.5, respectively, to the scRNA-seq data collected from Srrm2^+/-^ and WT cells, using the scmap label transfer method (Kiselev et al., 2018). As expected, when compared to early development cell references, the majority of WT cells are assigned to the earlier epiblast cells and only a small fraction is assigned to the later stage (93% to E4.5 and 6.7% to E5.5 epiblast; **Fig. 3G**). In contrast, the Srrm2^+/-^ cells are more frequently assigned to the later-stage epiblast E5.5, although the majority of cells are still assigned to the early epiblast cells of age E4.5 (76% to E4.5 and 20% to E5.5 epiblast). Furthermore, a larger fraction of Srrm2^+/-^ cells resembled committed cell types, such as trophectoderm, extra-embryonic ectoderm and primitive endoderm (3.8% in Srrm2^+/-^ and 0.3% in WT, **Fig. S3G**). When compared to the later stage embryos (E6.5-8.5), we observed that most of the WT (98%) and Srrm2^+/-^ cells (94%) were assigned to the epiblast (**Fig. 3H**). However, a larger subset of Srrm2^+/-^ cells (6.1%) compared to WT cells (2.4%) were transcriptionally most similar to differentiated cell types, without an evident cell-type preference (**Fig. S3H**). Taken together, these results suggest that Srrm2^+/-^ cells retain transcriptional profiles of early embryonic phenotypes, with a tendency to lose epiblast expression signatures.

### Transient *Srrm2* knockdown reveals alterations in specific splicing events

Gene expression alterations observed in the constitutive Srrm2^+/-^ heterozygous knockout ESC can comprise both direct effects of the SRRM2 protein, but also cellular responses due to genetic compensation response to the permanent depletion. To identify genes associated with the earliest cascade of deregulation driven by *Srrm2* dosage decrease, we used RNA interference (RNAi) to knockdown *Srrm2* in WT E14tg2a.4 cells for 24h and 72h, using as control small interfering RNA (siRNA) that target the firefly luciferase gene Gl2 (Srrm2-KD and Gl2-KD, respectively; **Fig. 4A**). qRT-PCR analyses confirmed depletion of *Srrm2* transcript to 50 and 40 % at 24h and 72h, respectively (**Fig. 4B**). RNAi-induced reduction in the dosage of *Srrm2* transcripts did not induce noticeable changes in the morphology of ESC colonies or an increased number of colony escapees (**Fig. S4A**), suggesting that conditional *Srrm2* depletion does not immediately interfere with pluripotency.

**Fig. 4.**
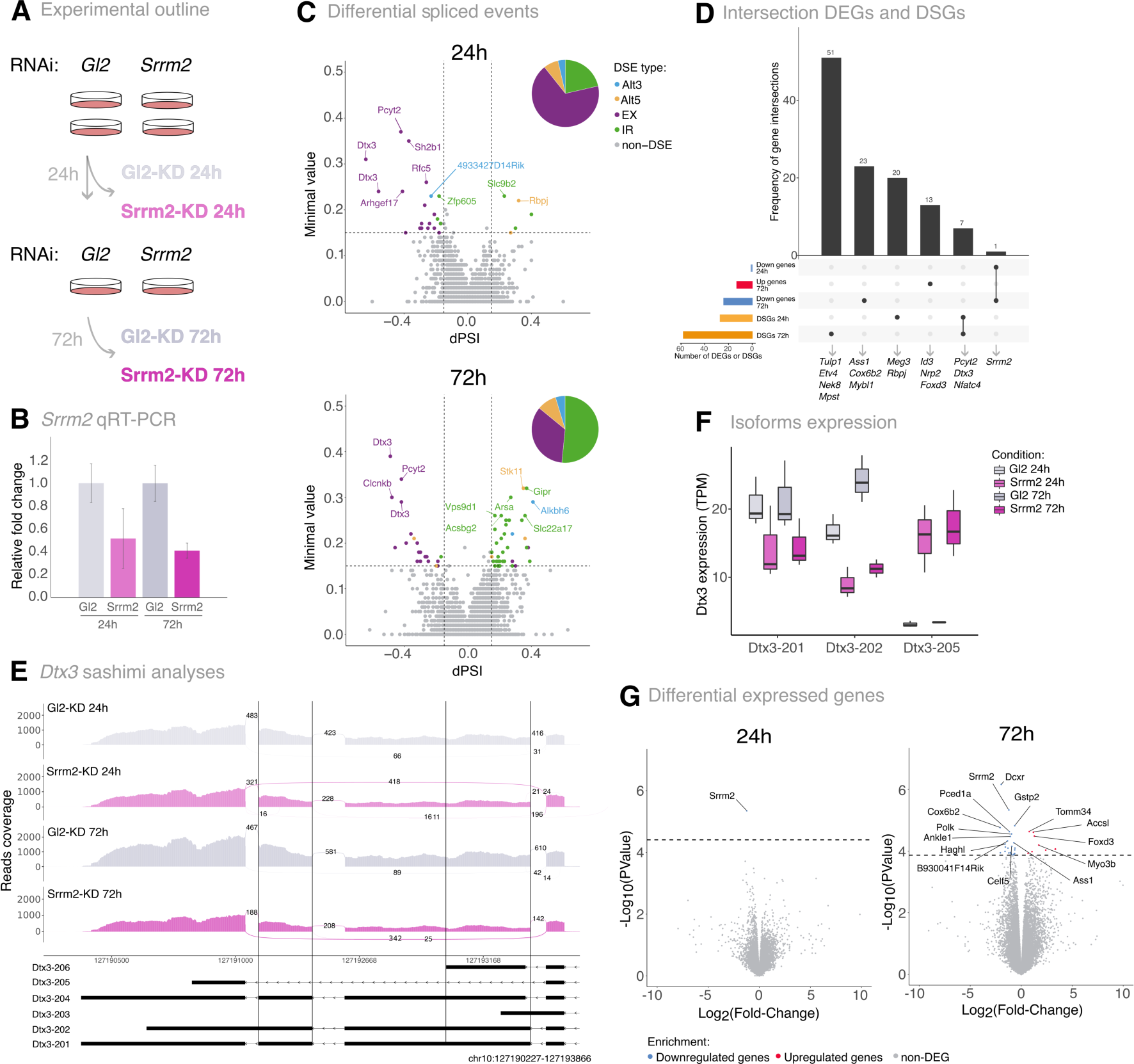
Transcriptome alterations upon *Srrm2* RNAi knockdown. A) Overview of the experimental outline. mESC were treated with RNAi targeting Gl2 luciferase gene (control) or *Srrm2*. Cells were harvested at 24h or submitted to another round of RNAi and harvested at 72h. RNA was extracted simultaneously. **B)** *Srrm2* expression levels. Total RNA levels of exon 11 of *Srrm2* were measured by RT-qPCR. Relative levels are normalized to *β-actin*. Mean and standard deviation are calculated from three replicates. **C)** Volcano plot showing the differentially spliced events upon *Srrm2* knockdown at 24h (top) and 72h (bottom). The magnitude of alterations is represented by the difference in Percent Spliced In (dPSI) and the minimal value. 324,859 and 303,695 splicing events were detected at 24h and 72h, respectively. 28 and 64 splicing events were considered as significantly differentially spliced, at 24h and 72h, respectively, using a cut-off of abs(dPSI) ≥ 0.15 and a minimal value ≥ 0.15. Spliced events with a dPSI < 0 correspond to exclusion events in *Srrm2* RNAi-treated cells, whereas dPSI > 0 correspond to inclusion events in *Srrm2* RNAi-treated cells. Categories of significant differential spliced events are represented by different colours: alternative 3’ splice site (Alt3, blue), alternative 5’ splice site (Atl5, orange), exon exclusion (EX, purple), intron retention (IR, green), no alteration (non-DSE, grey). The pie chart represents the proportion of significant differential spliced events. Gene labels represent the top 10 genes undergoing the differentially spliced events with the highest minimal value. **D)** Upset plot representing the comparison of differentially altered genes upon *Srrm2* knockdown at the expression and splicing levels for 24h and 72h. **E)** Sashimi plots displaying the differential splicing events for the *Dtx3* gene region (chr10:127190227-127193866) in control samples and Srrm2 knockdown samples at 24h and 72h. Splicing events are delimited with vertical black lines (*Dtx3*: chr10:127192609-127193357 and chr10:127191336-127191553). Numbers in the plot represent the average number of junction reads from three replicates. Isoforms that cover the represented genomic region are shown and identified as following: ENSMUST00000144322 (Dtx3-206), ENSMUST00000137151 (Dtx3-205), ENSMUST00000130855 (Dtx3-204), ENSMUST00000125254 (Dtx3-203), ENSMUST00000116229 (Dtx3-202), ENSMUST00000038217 (Dtx3-201). **F)** Expression levels (TPM) of *Dtx3* isoforms upon Gl2 (control) and *Srrm2* knockdown at 24h and 72h. Isoforms with expression levels above 5 TPM and encoding for proteins are displayed. **G)** Volcano plots showing the differentially expressed genes upon Srrm2 knockdown at 24h (left) and 72h (right). 19,461 genes were detected in the whole dataset, at 24h and 72h. 1 and 37 genes were considered as differentially expressed, at 24h and 72h, respectively, using an adjusted P-value < 0.05 as a cut-off (downregulated and upregulated genes are marked in blue and red, respectively).

To determine genome-wide changes in splicing events, we collected poly-adenylated RNA at 24h and 72h upon RNAi treatment in three independent biological replicates. We found significant differentially splicing events (DSEs) in 78 differentially spliced genes (DSGs), with 28 DSEs detected in 27 genes at 24h, and 64 DSEs detected in 58 genes at 72h (**Fig. 4C, Table S11-12**). The splicing alterations corresponded predominantly to exon-skipping events at 24h, similar to the type of splicing alterations identified in constitutive Srrm2^+/-^ cells (**Fig. 2D**), and were followed by an increase in intron retention events in the longer 72h exposure. All exon skipping events in the *Srrm2*-KD were characterized by exon exclusion without alterations in microexon splicing, as previously reported in other systems (Xu et al., 2022; Torres-Méndez et al., 2019; Cui et al., 2023).

GO analyses revealed no statistically significant enrichment in specific biological functions of the 78 DSGs, but a closer inspection revealed 22 genes associated with development, including *Pcyt2* (phosphate cytidylyltransferase 2), *Dtx3* (Deltex E3 Ubiquitin Ligase 3), transcription factors *Rpbj, Nfatc4*, *Etv4* and *Tulp1*, and long non-coding RNA *Meg3*. Interestingly, *Rbpj* is also a cluster 2 marker in the scRNA-seq analyses and is associated with exit from naive pluripotency to the formative state (Kalkan et al., 2019; Lackner et al., 2021). NFATC4 is a transcription factor required for cardiac development (Bushdid et al., 2003; Graef et al., 2001), which undergoes an exon skipping event that alters the relative abundance of protein-coding isoforms (**Table S13**). ETV4 expression can impair ectodermal commitment (Akagi et al., 2015), and also undergoes an exon-skipping event in *SRRM2*-depleted differentiated human cells (Cui et al., 2023). TULP1 is important for neuron and photoreceptor cell development (Jia et al., 2022), and *Meg3* transcripts sponge the microRNA *miR-423-5p* that in turn downregulates *Srf* expression (Cheng et al., 2020), which is upregulated in Srrm2^+/-^ cells.

Seven genes had common differential splicing events between 24h and 72h (**Fig. 4D**), with 8 common differential splicing events, all of which are exon cassette skipping (**Fig. S4B**). Among these genes, *Dtx3* and *Pcyt2* were also differentially spliced in Srrm2^+/-^ cells, with two and one exon skipping events, respectively (**Fig. 2C**), showing that their splicing deregulation occurs upon acute *Srrm2* depletion and prevails in an adapted heterozygous knockout ESC. DTX3 can act both as an activator or repressor of NOTCH signalling, a key pathway that regulates cell fate determination (Ding et al., 2020; Wang et al., 2021b). *Pcyt2* is an embryonic-lethal gene which encodes a protein involved in phospholipid synthesis (Fullerton et al., 2007). Visual inspection of the density of RNA-seq reads along splice junctions, using sashimi plots, confirmed that the skipped exons have a lower proportion of coverage of sequencing reads at both genes, in Srrm2-KD cells at both 24h and 72h (**Fig. 4F; Fig. S4C**).

To further explore the impact of the exon skipping in *Dtx3* and *Pcyt2*, we compared the expression levels of their annotated protein-coding isoforms. At both 24 and 72h, we found a decrease in the two highest expressed *Dtx3* protein-coding isoforms (Dtx3-201 and Dtx3-202) accompanied by a strong upregulation of another protein-coding isoform (Dtx3-205) (**Fig. 4F**). Notably, Dtx3-205 encodes for a DTX3 protein that lacks a RING domain present in Dtx3-201 and Dtx3-202, which is responsible for ubiquitination transfer. For *Pcyt2*, the highest expressed isoform, which encodes the canonical PCYT2 (PCYT2α) protein isoform, did not show notable expression differences, whereas a Pcyt2 protein-coding isoform that lacks exon 7 (Pcyt2β) was more highly expressed at both 24h and 72h (**Fig. S4D**). The corresponding protein isoform has the same functional domains as Pcyt2α, but differs in catalytic and kinetic activities and is expressed in a tissue-specific manner (Tie and Bakovic, 2007). *Pcyt2*α is more prominently expressed than *Pcyt2*β in most mouse tissues, except in muscle and heart tissues where their expression is similar. The same splicing alteration in PCYT2 was reported after SRRM2 depletion in the human hepatoma cell line HepG2 (Cui et al., 2023).

### Transient *Srrm2* depletion in ESC leads to mild changes in genome-wide gene expression

Finally, we investigated gene expression changes upon *Srrm2* RNAi treatment. At 24h, only *Srrm2* was significantly downregulated, while 24 genes were down-regulated at 72h, including *Srrm2*, and 13 were upregulated (**Fig. 4G; Tables S14-15**). Among the 78 differentially spliced genes in Srrm2- KD, none is differentially expressed at 24h or 72h (**Fig. 4D**), suggesting that the splicing alterations at specific genes related with *Srrm2* dosage decrease precede the more global gene expression changes detected in constitutive *Srrm2* heterozygosity.

Within the 37 differential expressed genes detected in Srrm2-KD cells, we found embryo development-related and transcription factor genes, such as *Id3*, *Mybl1*, *Nrp2*, *Fgf11* and *Foxd3* (**Fig. 4G**), with *Id3, Mybl1 and also Cox6b2*, being differentially expressed in the same direction in Srrm2^+/-^ cells (**Fig. S4E**). Out of the 37 genes found differentially expressed upon *Srrm2* knockdown, only 16 were differentially expressed in the constitutive Srrm2^+/-^ cells, and only 9 were differentially expressed in the same direction (**Fig. S4E**). These results suggest that the imbalance in cell identity found upon constitutive *Srrm2* dosage reduction in Srrm2^+/-^ cells, likely results from a few direct effects of SRRM2 likely through splicing deregulation, which lead to a cascade of indirect effects on stemness and cell lineage regulatory networks.

To zoom in further on the genes that are deregulated in both constitutive and RNAi depletion of *Srrm2*, we compared the expression of the Srrm2-KD DEGs with the earliest gene expression alterations of Srrm2^+/-^ cells (**Fig. S4F**). Although the correlation of expression was very low (R = 0.3, P-value = 0.12), we found that the genes that are differentially expressed after RNAi also tend to change expression in the same direction in the most pluripotent states of Srrm2^+/-^ cells detected by scRNA-seq, clusters 1 and 2. Amongst these, we find *Cox6b2*, a gene marker of cluster 1, and significantly downregulated in cluster 2, and at 72h upon *Srrm2* RNAi treatment (**Fig. 4G, Fig. S4F- G**). *Ass1*, which encodes for a protein that catalyses the penultimate step of the arginine biosynthetic pathway, is also a marker gene of cluster 1, is found significantly downregulated at 72h in Srrm2- KD cells, and its expression decreases from cluster 1 to 2, although not significantly. In contrast, the myogenesis-related transcription factor *Id3* is upregulated 72h after *Srrm2* depletion and it is highly expressed in cluster-4 state cells, corresponding to the most differentiated group of ESC within the cultures.

Taken together, our results show that transient depletion of *Srrm2* leads to exon cassette exclusion and intron retention splicing alterations in specific genes, including development-related genes.

These specific splicing alterations precede gene expression alterations, suggesting that a small number of specific SRRM2-dependent splicing defects drive the gene expression alterations that ultimately lead to an impairment in the stemness of ESC upon *Srrm2* depletion.

## Discussion

In this work, we investigated the roles of *Srrm2* dosage in stemness through the modulation of splicing and gene expression networks, using *Srrm2* heterozygous knockout or transient knockdown in mouse ESC. The RNAi knockdown revealed that the most acute effects of *Srrm2* half dosage are alternative splicing defects in a small number of genes with roles in gene and metabolism regulation, without immediate changes in gene expression levels. In contrast, single-cell transcriptomic analyses showed that the constitutive half dosage of *Srrm2* in ESC gives rise to heterogeneous cultures reliably composed of cells in different transcriptional and stemness states. Embryonic stem cells have the unique capacity to differentiate into a wide range of specialized cell types (pluripotency) and to perpetually propagate in culture while retaining their undifferentiated state through self-renewal (Martin, 1981; Evans and Kaufman, 1981; Thomson et al., 1998; Huang et al., 2015). Our study reveals that a persistent reduction in *Srrm2* transcripts in stem cells reduces the competency to maintain stemness and results in the emergence of a mixed cell population with distinct phenotypes, which include defective colony formation and altered expression of pluripotency markers. Srrm2^+/-^ cells significantly overexpress genes related to development and cytoskeleton, whereas the group of downregulated genes are enriched in genes associated with the LIF pathway, crucial for ESC self-renewal (Smith et al., 1988; Williams et al., 1988; Huang et al., 2015). Notably, LIF withdrawal or the direct inhibition of downstream effectors leads to differentiation of mouse ESC into morphologically mixed cell populations, similar to what we observe in Srrm2*^+/-^* population (Niwa et al., 1998). We identify *Srrm2* dosage as a key player in balancing self-renewal and differentiation processes, to ensure appropriate proportions of stem cell niches.

Single-cell RNA-seq analysis revealed distinct cell states among Srrm2^+/-^ cells. While the majority of Srrm2^+/-^ cells retain pluripotency expression signatures, namely *Pou5f1*, *Sox2*, *Nanog*, *Klf4*, their pluripotency state is markedly different from the most abundant pluripotent state in WT cells. Mouse ESC cultured in the presence of differentiation-restricting inhibitors, such as LIF, are *in vitro* counterparts of the naive preimplantation epiblast E4.5. EpiLCs, derived from mESC, resemble the early post-implantation epiblast (E5.5-6.5) and exhibit features of the formative pluripotent state which precedes the induction for multi-lineage specialization (Hayashi et al., 2011; Rossant and Tam, 2017; Kurimoto et al., 2015; Yang et al., 2019). Compared to the WT condition, the population of Srrm2^+/-^ ESC contains a higher proportion of cells resembling E5.5 stage cells, and we found that most Srrm2^+/-^ ESC exist in an intermediate state between the naive and formative pluripotency. This intermediate state is characterized by a reduction in the expression of *Nanog* and *Klf4*, and overexpression of the DNA methyltransferases, *Dnmt3a* and *Dnmt3b* and several ribosomal protein genes, such as *Rps21*, *Rps29*, *Rpl29*, *Rpl36*. Moreover, transient Srrm2 depletion by RNAi has a rapid effect on the expression of metabolic-related genes. In particular, we found misregulation of *Cox6b2*, involved in mitochondria oxidative phosphorylation, and *Ass1*, a component of the arginine pathway important in protein synthesis. Metabolic cues are tightly regulated during pluripotency and differentiation, and proper regulation of ribosomal gene expression is crucial for the requirements of different cell states. This highlights a possible role of *Srrm2*, and more specifically its dosage, as an early modulator of gene expression networks, but also in controlling metabolic pathways, both key to balancing a coordinated and dynamic progression of pluripotent states during embryo development.

The modulation of transcription factor expression cascades, and the putative alteration of the epigenetic and metabolic landscape in Srrm2^+/-^ cells, suggests that *Srrm2* depletion leads to a partial dismantling of stem regulatory networks, promoting the onset of cell lineage commitment, into a state where the majority of cells are beyond their naive pluripotency features but are still not committed to a particular lineage. Alternatively, it is possible that the intermediate state of Srrm2^+/-^ cells between naive and formative pluripotency does not exist *in vivo* in embryos, or in standard cultures of ESC. Regardless, our study suggests that *Srrm2* levels play important roles in coordinating the transitions or the temporal dynamics between naive and formative pluripotent stem cells, and the robustness of their stemness. Further studies may help elucidate possible roles of Srrm2-mediated mechanisms in promoting fluctuations of pluripotent states that provide windows of opportunities for cell commitment.

Srrm2^+/-^ cells express cell-type-specific markers and transcription factors important for mesoderm or cardiac tissues. Importantly, upon *Srrm2* RNAi, we also observed misregulation of genes associated with mesoderm and angiogenesis, such as *Id3*, *Myo3b* and *Nrp2*, before detectable changes in pluripotency regulation genes. A stable and continuous depletion of *Srrm2* in Srrm2^+/-^ mESC leads to a significant enrichment of the *Srf* transcription factor, a master regulator of cardiac development, and its target genes, especially in the most differentiated cells. SRF activation is driven by cell mechanic cues such as extracellular physical or chemical signals that are transduced by actin cytoskeleton remodelling signalling cascades coordinated with activation of myocardin-related transcription factors (Olson and Nordheim, 2010; Posern and Treisman, 2006; Xiong et al., 2022).

Cytoskeletal and mechanical cues play key roles in regulating pluripotency by modulating the transcriptome of mouse ESC in response to external signalling cues (David et al., 2019; Heo et al., 2017; Putra et al., 2023). The misregulation of cytoskeletal-related genes in Srrm2^+/-^ cells, such as Krt8 and Krt18, could potentially identify important players in controlling cell mechanical cues, which may be interconnected with alterations in SRF-related pathways. Transient depletion of *Srrm2* led to splicing alterations in *Meg3* lncRNA, which indirectly regulates *Srf* transcript levels through its sponging functions. Alternative splicing of *Meg3* could lead to alterations in its three-dimensional folding, impacting its sponging abilities, and potentially explaining the increase in SRF targets.

Further studies are required to better understand whether the alterations in Meg3 isoforms, cytoskeleton and SRF-pathway drive or result from the loss of stemness in Srrm2^+/-^ ESC.

While Srrm2^+/-^ cells show extensive gene expression alterations with few splicing alterations, transient *Srrm2* depletion leads to minor expression changes and reveals that splicing misregulation, mostly involving exon skipping and intron retention, as previously shown in differentiated cells (Cui et al., 2023; Xu et al., 2022), occurs before gene expression effects. Notably, *Dtx3* and *Pcyt2* exhibit differential splicing events in protein-coding isoforms both in acute and constitutive *Srrm2* dosage reduction, indicating that they may play pivotal roles in triggering the cascade of *Srrm2*-associated phenotypes. DTX3 is an ubiquitin ligase with a key role in cell fate determination and is a regulator of Notch signalling, which governs numerous processes during embryo development. PCYT2 is an enzyme involved in the Kennedy pathway of phospholipid synthesis, and its roles in development are still not known. These findings suggest that *Srrm2*-dependent splicing of specific genes, including *Dtx3* and *Pcyt2*, as a likely mechanism driving subsequent changes in gene expression and ultimately impairing pluripotency.

Further research is needed to provide insights into the molecular mechanisms underlying the effects of *Srrm2* dosage reduction in pluripotency maintenance and early differentiation. Recent studies performed in differentiated cells have placed SRRM2 in the spotlight of splicing speckles composition and biogenesis (Ilik et al., 2020; Xu et al., 2022; Hu et al., 2019). For a better understanding of SRRM2-mediated mechanisms, it will be important to disentangle its role in alternative splicing, splicing speckle formation and genome organization in the context of pluripotency and development.

Taken together, our study sheds light on the roles of splicing factors in the intricate interplay between alternative splicing and gene expression networks that govern early developmental processes. *Srrm2* emerges as a key regulator of pluripotency and early development potentially through its dosage-dependent regulation of specific splicing events and its ability to balance unscheduled gene expression alterations. Furthermore, the identification of early and long-lasting *Srrm2*-mediated splicing and expression defects at specific genes prompts further investigation of their roles in biological and medical contexts, opening opportunities for exploring new therapeutic strategies targeting *Srrm2* and its downstream effectors in developmental and cancer diseases.

## Materials and methods

### *Srrm2* expression analysis during *in vivo* mouse development

*Srrm2* expression levels were assessed during E3.5-5.5 mice embryo development using publicly available data (Nowotschin et al., 2019), and the force-directed layouts plots in Fig. 1A were produced using the interactive data website www.endoderm-explorer.com. Gene expression color per cell represents the post-imputation with the MAGIC algorithm (van Dijk et al., 2018) following normalization and log transformation. *Srrm2* expression levels during E10-P63 mice embryo development were assessed using publicly available data (Cardoso-Moreira et al., 2019). The data used to produce Fig. 1B was obtained from the interactive app https://apps.kaessmannlab.org/evodevoapp/.

### mESC lines

Srrm2^+/-^ cells (CG0630) were generated by a gene trap-derived truncation in the *Srrm2* gene by the Sanger Institute Gene Trap Resource (SIGTR), Cambridge, UK, using mouse ESC lines clone E14tg2a.4, as a recipient. Srrm2^+/-^ and parental WT E14tg2a.4 ESC were obtained from the Mutant Mouse Resource Center (MMRRC) at the University of California at Davis, a National Institutes of Health (NIH) funded strain repository. MMRRC screening in Srrm2^+/-^ cells reported *Mycoplasma spp.* and mouse Parvo viruses free, and a chromosome count of 40 chromosomes. E14tg2a.4 WT cells were expanded and tested negative for *Mycoplasma spp.*, upon screening according to the manufacturer’s instructions (AppliChem, A3744). E14tg2a.4 WT and Srrm2^+/-^ cells were expanded and cryopreserved at three different passage numbers (WT: passage 19-21, Srrm2^+/-^: passage 4-7) in three independent batches, that were further thawed and treated as biological replicates. Bulk sequencing datasets produced from E14tg2a.4 WT and Srrm2^+/-^ were mapped and tested negative for *Mycoplasma spp*.

### mESC culture

mESC were grown as previously described (Beagrie et al., 2017). Briefly, cells were grown at 37°C, 5% (v/v) CO2, on gelatine-coated (0.1% v/v, Sigma-Aldrish, cat# G1393) Nunc T25 flasks in Gibco Glasgow’s MEM (Invitrogen, 21710-082), supplemented with 10% (v/v) heat-inactivated FCS (Invitrogen, 10270-106, batch number 41F8126K), 2,000 units ml^−1^ LIF (Millipore, ESG1107), 0.1 mM β-mercaptoethanol (Invitrogen, 31350-010), 2 mM L-glutamine (Invitrogen, 25030-024), 1 mM sodium pyruvate (Invitrogen, 11360-070), 1% penicillin–streptomycin (Invitrogen, 15140- 122) and 1% non-essential amino acids (Invitrogen, 11140-035). The cell culture medium was changed every day and cells were split every second day. Before sample collection, cells were grown for 24h, 48h or 72h in serum-free ESGRO Complete Clonal Grade medium (Merck, SF001-500P), supplemented with 1,000 units ml^−1^ LIF, on 0.1% gelatine-coated (Sigma, G1393-100 ml, 0.1% v/v) Nunc dishes, with a daily medium change. Cultures were visually inspected and brightfield images were acquired in an Olympus IX53 microscope.

### Alkaline Phosphatase assay

WT and Srrm2^+/-^ E14tg2a.4 cells were seeded at a clonal density of 8.5x10^3^ cells/cm^2^, and grown for 72h under normal conditions. The Alkaline Phosphatase assay was carried out according to the manufacturer’s instructions (Sigma-Aldrich, 86R-1KT). Alkaline phosphatase activity is highest in stem cells, and it reduces as cells differentiate into other cell types, thus AP activity is commonly used as a means of determining the ability of cells to differentiate. Cells with higher AP activity show higher Hematoxylin staining. AP staining was visually inspected and brightfield images were acquired in an Olympus IX53 microscope.

### *Srrm2* RNAi knockdown in mESC

WT E14tg2a.4 ESC were transfected using OptiMEM (Invitrogen, 31985062), Lipofectamine RNAiMAX (Invitrogen, 13778075) and 50 nM synthetic siRNA duplexes (Eurogentec) targeting either *Srrm2* or firefly luciferase (Gl2), which was used as a control. RNA interference was done according to the manufacturer’s instructions. Briefly, cells were seeded and reverse-transfected at a density of 3.1x10^4^ cells/cm^2^. 24h after the first transfection, cells were harvested or re-transfected with the same siRNA duplexes and transfection reagents and harvested 72h after seeding. The siRNA sequences are reported from 5’ to 3’: Gl2 sense CGUACGCGGAAUACUUCGA, Gl2 antisense UCGAAGUAUUCCGCGUACG, *Srrm2* E8/9 sense GAAGAAGCACAGGUCAGAA, *Srrm2* E8/9 antisense UUCUGACCUGUGCUUCUUC, *Srrm2* E13 sense CGUGGAAGAUCUCACUCUA, *Srrm2* E13 antisense UAGAGUGAGAUCUUCCACG.

### RNA isolation

For total RNA-seq and qRT-PCR, Srrm2^+/-^ and WT ESC were seeded (5.2x10^4^ cells/cm^2^) and grown for 48h to 70-80% confluency. Total RNA was isolated with TRIzol (Invitrogen, 15596-026) following the manufacturer’s instructions. Lysates were either processed immediately for qRT-PCR, or snap-frozen in liquid nitrogen and stored at -80°C until further processing. DNA was degraded with TURBO DNase (Invitrogen, AM1907), according to the manufacturer’s protocol. Purified RNA was either stored at -80°C or processed immediately.

### cDNA synthesis and real-time quantitative PCR

Purified RNA (1 μg) was reverse transcribed into cDNA using 10 U/μl of Superscript II Reverse Transcriptase (Invitrogen, 18064-022), 50 ng of random primers (Promega, C118A), 0.5 mM dNTPs (NEB, N0447L), 5 mM RNAseOUT (Invitrogen, 10777-019) in a 20 μl solution with 10 mM DTT and Superscript buffer, according with the manufacturer’s instructions. RNA-DNA hybrids were removed with 2 U of RNase H (NEB, M0297S). The synthesized DNA was diluted 1:4 and 1 μl was used for qRT-PCR, using SensiMix™ SYBR^®^ No-ROX (Meridian Bioscience, QT650-05), 250 nM of each primer in a final volume of 20 μl. The primers sequences are reported from 5’ to 3’: *Srrm2* Exon 11 FW AGTCTCTCGTAGAAGCCGGT, *Srrm2* Exon 11 RV CTTCTGCGTCTTGGTGGAGT, *Actb* FW TCTTTGCAGCTCCTTCGTTG and *Actb* RV ACGATGGAGGGGAATACAGC. *Srrm2* expression was measured from 3 biological replicates per condition, and were assessed according to the 2–ΔΔCT method (Livak and Schmittgen, 2001), and normalized to *Actb*. The relative RNA expression for each condition was estimated by calculating the ΔCT average per replicate. The mean and standard deviation (S.D.) were then determined from the replicates.

### Immunofluorescence

Srrm2^+/-^ and E14tg2a.4 WT cells were seeded (1x10^5^ cells/cm^2^) on ethanol-washed and autoclaved glass 0.1 % gelatin-coated coverslips and grown under regular conditions for 1 day. Cells were rinsed and fixed in 4% EM-grade PFA (Alfa Aesar, 43368) in 125 mM HEPES-NaOH pH 7.6, for 30 min at room temperature. Aldehydes were quenched (10 min) with 20 mM glycine in PBS. Cells were permeabilized (10 min) in 0.5% Triton X-100 in PBS (w/v). Cells were washed and incubated (3x over 1h) in blocking solution (1% BSA, 0.2% fish skin gelatin, 0.1% casein, in PBS, pH 7.8).

Primary antibodies were diluted in blocking solution and incubated overnight at 4°C in a humid chamber. Cells were washed (3x over 1h) in blocking solution, before incubation with secondary antibody diluted in blocking solution for 1h at RT. After incubation, cells were washed in PBS 3 times for 1 h. DNA was stained with 0.5 µg/ml DAPI (Sigma-Aldrich, D9542) in PBS for 2 min. Cells were washed 3 times in PBS and mounted in Vectashield (Vector Laboratories, H-1800) before imaging. The antibodies used are listed in Table S16.

### Confocal imaging

Immunofluorescence images were acquired with a Leica TCS SP8-STED confocal microscope (Leica Microsystems DMI6000B-CS) using Leica Application Suite X v.3.5.5.19976 and an HC PL APO CS2 63x/1.40 oil objective, with a pinhole equivalent to 1 Airy disk. Images were acquired using 405 nm, 488 nm, 555 nm, 633 nm and 680 nm excitation using a long-pass filter at 1024 x 1024 pixel resolution (pixel size 180 x 180 nm). For intensity signal comparison, representative images from the same experiment set were collected without intensity signal saturation, in the same session with the same settings. Images were post-processed using Fiji software (version 2.9.0/1.53t; Schindelin et al., 2012; Linkert et al., 2010), and comparative images were contrast-stretched using the same parameters.

### Western blot

E14tg2a.4 WT and Srrm2^+/-^ cells were seeded (5.2x10^4^ cells/cm^2^) and grown for 48h to 70-80% confluency. The organic phase was extracted following TRIzol protocol (Invitrogen, 15596-026), according to the manufacturer’s instructions. Proteins were precipitated with isopropanol and washed with 0.3 M of guanidine hydrochloride in 95% ethanol for 20 min. Protein pellets were resuspended in 1% SDS and protein concentration was measured using the Bradford assay kit (Thermo Scientific, 1863028), according to the manufacturer’s instructions. 10 µg of protein extracts were denatured in 5x Laemmli-buffer (125 mM Tris-HCl pH 6.8, 10% SDS, 50% glycerol, 0.5% bromophenol blue, 0.2% ꞵ-mercaptoethanol) for 5 min at 95°C. Protein lysates were loaded on 10% polyacrylamide gels and resolved at 120 V for 2h at RT. Proteins were transferred to a PVDF membrane using a Mini Trans-Blot® Cell (Bio-Rad Laboratories) at 100V for 2h, at 4°C. The membrane was blocked in 5% skimmed milk for at least 1 h. Primary antibodies incubation was performed overnight at 4°C and secondary antibodies were incubated for 2h. Chemiluminescent substrate was added according to the manufacturer’s instructions (Amersham ECL Prime Western Blotting Detection Reagent, Cytiva #RPN2232) and was detected with an Amersham Imager 600 (GE Healthcare, 10600023). The antibodies used in this study are listed in Table S16.

### Flow cytometry

Srrm2^+/-^ and E14tg2a.4 WT cells were grown under regular conditions for 48h. Triplicate wells (technical replicates) were each harvested at three different passages using TrypLE (Gibco). Cells were fixed and permeabilized using the permeabilizing buffers from the FoxP3 staining buffer kit (Miltenyi; 130-093-142), and stained with conjugated antibodies (Table S16). Protein expression was analyzed using a MACSQuant VYB flow cytometer (Miltenyi). Gating and plots were generated using FlowJo 10.

### Bulk RNA-seq library preparation and sequencing

Purified RNA was produced in three biological replicates from WT and Srrm2^+/-^ E14tg2a.4 cells, or from RNAi-treated E14tg2a.4 WT cells, and tested for RNA quality using the 2100 Bioanalyzer Agilent system with the Agilent RNA 6000 Nano assay (Agilent Technologies, 5067-1511). RNA spike-ins (Invitrogen, 4456740) were added to each purified RNA sample following the manufacturer’s recommendations. Total RNA libraries were produced from 600 ng of RNA using the TruSeq Stranded total RNA library preparation kit (Illumina, 20020596) according to the manufacturer’s protocol. PolyA+ RNA-seq libraries were produced from 1 µg of RNA using the TruSeq® Stranded mRNA Library Prep (Illumina, 20020594) kit, according to the manufacturer’s instructions. Libraries were analysed on the 2100 Bioanalyzer Agilent system using the Agilent High Sensitivity DNA kit (Agilent Technologies, 5067-4626). DNA concentrations were measured by Qubit Quant IT kit (Invitrogen, Q10212) according to the manufacturer’s protocol. Total RNA libraries were paired-end sequenced using the NextSeq 500/550 system at 75 bp length, according to the manufacturer’s protocol. Sequencing depth ranged from 78-99 million reads in WT and Srrm2^+/-^ replicates. PolyA^+^ RNA libraries were paired-end sequenced using the NovaSeq 6000 system at 100- bp length, according to the manufacturer’s protocol. Sequencing depth ranged from 91-251 million reads per replicate.

### Total and polyA+ RNA-seq data analyses

#### Gene expression

Total and polyA expression levels from RNA-seq profiles were obtained using Kallisto v0.46.1 (Bray et al., 2016), where reads were pseudo-aligned to the mouse gencode transcriptome release M24. Differential expression analysis was conducted using edgeR (version 3.38.4). Briefly, sample comparison was performed by filtering for low expressed genes, applying a quasi-likelihood method (glmQLFTest function), and selecting significantly differentially expressed genes with FDR adjusted P-values lower than 0.05, as implemented in the edgeR package. For polyA+ RNA-seq, an additional step of batch effect correction for the processing date of the samples was performed indicating the RNA processing day as a covariate in the design matrix. To assess the accuracy of the transcriptome alteration, we analysed the correlation between the observed and expected fold changes of the spike-ins (total RNA: R = 0.97, P-value<2.2x10^-6^; 24h and 72h polyA+ RNA: R = 0.98, P-value<2.2x10^-6^).

#### Gene Set Enrichment and over-representation analyses

Gene set enrichment analyses (GSEA) and over-representation analysis (ORA) were performed with WEB-based GEne SeT AnaLysis Toolkit (WebGestalt 2019), using the default parameters and weighted set to reduce redundancy of the gene sets in the enrichment result (Zhang et al., 2005; Wang et al., 2013; Wang et al., 2017; Liao et al., 2019). For GSEA all the detected genes were used as an input and ranked according to their P-value and the sign of the fold change, as previously described (Reimand et al., 2019). GSEA was performed for the default functional databases: biological processes (Gene Ontology) and predicted transcription factor targets of networks (Gene Transcription Regulation Database, GTRD) (Kolmykov et al., 2020). Furthermore, external databases were used as functional databases in WebGestalt to perform GSEA and ORA. To assess commitment to a specific cell lineage using signatures previously identified by single-cell transcriptome profiles of mouse embryos (Cao et al., 2019) and retrieved from MSigDB’s hallmark collection (https://www.gsea-msigdb.org/; Liberzon et al., 2015). To identify the transcription factors necessary to induce cell-type reprogramming, we used the publicly available Mogrify (Rackham et al., 2016) gene lists (Ferrai et al., 2017) as a functional database to perform an ORA, providing Srrm2^+/-^ upregulated genes as input and all detected genes as background. Finally, the list of SRF targets predicted by the GTRD was downloaded from the Human Molecular Signatures Database (MSigDB), C3 subcollection TFT: Transcription factor targets and converted into *mus musculus*, using the Bioconductor BiomaRt R package and the *Mus musculus* transcription factors list was obtained from the AnimalTFDB 3.0 database (Hu et al., 2018).

#### Splicing alterations

For AS event-level analysis, raw reads were mapped to previously annotated AS events for the mouse reference assembly (mm10 vastdb.mm2.23.06.20) from VastDB (VastDB v2 released). Abundance of AS events in PSI values was estimated using Vast-tools (version 2.5.1) align and combine commands (Irimia et al., 2014; Tapial et al., 2017). Vast-tools quantifies exon and microexon skipping, intron retention, and alternative donor and acceptor site choices. For further analyses, we filtered splicing events that had at least 2 replicates per condition with a minimum read coverage of LOW. To calculate differential AS between samples, we used Vast-tools *diff* function. Vast *diff* uses Bayesian inference followed by differential analysis to calculate percent spliced inclusion variations (dPSI) and minimum value difference (MV) between two samples for each AS event. We selected, as significantly differentially spliced, events with an absolute dPSI ≥ 0.15 and a MV ≥ 0.1 or 0.15 (total and polyA+ RNA, respectively) at a 95% confidence interval. Significant events were individually assessed for the quality of the PSI distributions in both of the compared conditions. For visualization of splicing events, read alignments from 3 replicates were merged per condition and visualized in the form of sashimi plots, using ggsashimi tool v1.1.5 (Garrido-Martín et al., 2018). We applied a threshold of 10 minimum reads to support a junction in sashimi plots.

Expression levels for each individual isoform were estimated and normalized in Transcripts per Million (TPM) with Kallisto, as described above.

### Single-cell mRNA library preparation

Two independent replicates of Srrm2^+/-^ and E14tg2a.4 WT cells were grown for 48h under regular conditions. Cells were resuspended in 0.04% BSA in PBS and the suspension was loaded on a 40 µm Flowmi™ Tip Strainer to remove cell debris and clumps. The cell concentration and viability of the single-cell suspension were determined using a Countess® II Automated Cell Counter (Thermo Fisher Scientific, AMQAX1000), following the manufacturer’s instructions. Cell viability ranged between 86-96%. Cells were resuspended into 10^6^ cells/ml in 0.04% BSA in PBS. Single-cell mRNA-seq libraries were prepared with the Chromium Single Cell 3’ v3 chemistry Kit (10x Genomics), according to the manufacturer’s instructions. Briefly, a droplet emulsion was generated in a microfluidic chip followed by barcoded cDNA generation inside these droplets. Purified and amplified cDNA was then subjected to library preparation, and sequenced on a NovaSeq 6000 instrument (Illumina).

### Single-cell RNA-seq data analyses

#### Mapping, filtering, pre-processing, data integration, clustering and cell cycle assignment

The four 10x Genomics scRNA-seq libraries were simultaneously mapped to the mouse genome (refdata-gex-mm10-2020-A version provided by cellranger) using the *cellranger count* function from 10x Genomics Cellranger software (version cellranger-6.1.2) with default parameters, except - expect-cells 10,000 (total number of cells targeted per experiment). The four 10x Genomics scRNA-seq libraries were aggregated on one unique count matrix using *cellranger aggr* function from 10X Genomics Cellranger software (version cellranger-6.1.2) with default parameters. The analysis of the single-cell data was performed using Seurat version 4.1.0. To remove low-quality cells, we applied sample-specific thresholds for both number of UMIs and number of features, by subtracting one standard deviation from the mean value across all cells of UMIs or features, respectively. Cells that did not pass both of these two thresholds were removed from the analyses. Additionally, cells having higher than 10% of mitochondrial reads were also excluded.

Single-cell datasets were normalized using the *LogNormalize* method, with a scale factor of 10000, and the top 2000 variable genes were calculated using the *vst* method. Lastly, the data were scaled using the *ScaleData* function from Seurat, using all genes as input.

PCA analysis was performed by running the *RunPCA* function from Seurat with default parameters, and it was followed by the integration of all samples using Harmony (version 1.0). The batch-corrected data was subjected to uniform manifold approximation and projection (UMAP) using the first 10 dimensions. To explore distinct biologically meaningful groupings, we proceed with clustering by running *FindNeighbors* (first 20 dimensions) and *FindClusters* (0.2 resolution) from the Seurat package. To estimate the cell cycle phase of each cell in our study, we used the *project_cycle_space* and *estimate_cycle_position* from the tricycle package (version 1.2.1). For simplicity, numerical values were converted to categorical labels following the developer’s recommendations: cells with a tricycle position between 0.5pi and pi were classified as “S” phase, those between pi and 1.75pi as “G2M” phase, those greater than or equal to 1.75pi or less than 0.25pi as “G1/G0” phase, and the rest as “NA”.

#### Differential expression analyses and over-representation analyses

To identify cluster-specific marker genes that may characterize all the different cellular populations, we performed differential gene expression analysis using the *FindAllMarkers* function in Seurat.

This analysis was conducted using the Wilcoxon rank-sum test and considering only positively regulated genes detected in at least 10% of cells within each cluster with a minimum fold change of 0.25. We excluded clusters that were composed of less than 2% of the total number of cells in the dataset, specifically clusters 5 and 6. Analyses of the expression of pluripotency markers showed a progressive reduction in its levels along the identified clusters (Fig. 3, Fig. S3). This suggests that the UMAP projection captures a progressive state that underlies a loss of stemness from cluster 1 to cluster 4 and that a trajectory analysis would not provide additional insight into the cellular dynamics of our system. Pairwise differential expression analysis between cluster 2 (mostly composed of Srrm2^+/-^ cells, making up 95% of the cluster) and cluster 1 (mostly containing WT cells, making up 94% of the cluster) was performed using FindMarkers function (Seurat). Genes with Log2 Fold change > 0 represent upregulated genes in cluster 2, whereas genes with Log2 Fold change < 0 are upregulated in cluster 1. Differentially expressed genes were expressed in at least 50% of the cells in at least one of the two clusters being compared (min.pct = 0.5).

To investigate the pluripotency state of the single cells, we assessed the expression of gene markers, as previously described (Wang et al., 2021a).Over-representation analysis for biological terms (http://geneontology.org/) and phenotype terms (https://www.informatics.jax.org/vocab/mp_ontology/) were performed with WEB-based GEne SeT AnaLysis Toolkit (WebGestalt 2019), using the default parameters and weighted set to reduce redundancy of the gene sets in the enrichment result (Zhang et al., 2005; Wang et al., 2013; Wang et al., 2017; Liao et al., 2019). Gene markers were provided as input against a gene universe of all genes expressed in at least 10 of the cells that were included in the analysis. The top 5 terms with a higher Normalized enriched Score (NES) for each cluster were selected to produce the heatmaps shown in Fig. 3F and Fig. S3E.

#### Single-cell labelling transfer

Mice *in vivo* wild-type scRNA-seq data and cell type annotations from E3.5-5.5 and E6.5-8.5 development stages were acquired from GSE123046 (Nowotschin et al., 2019) and GSE137337 (Chan et al., 2019; Grosswendt et al., 2020; Haggerty et al., 2021) repositories, respectively. Data normalization was conducted using the Seurat package’s *NormalizeData* function (v. 4.3.0.1). For the E3.5-5.5 reference dataset, ICM cells were identified and annotated de-novo based on increased and specific expression of key pluripotency genes (*Pou5f1, Sox2*, *Nanog*, *Klf2*, *Klf5*). The most similar *in vivo* cell types were identified for wild type and Srrm2^+/-^ cells using the *scmapCluster* function (Scmap v. 1.22.3 using 2000 highly variable features and an assignment threshold of 0.5) of the Scmap cell-cluster label transfer method (Kiselev et al., 2018).

### Manuscript preparation

R version 4.2.1 (2022-06-23) and Inkscape version 1.2.1 (https://inkscape.org/) were used for data visualization and figure preparation, respectively.

### Data availability

Datasets produced in this study can be found in the GEO repository under the accession number GSE243433, which includes raw fastq files and raw count matrices for the RNA-seq bulk datasets, and fastq files and matrix processed files for single-cell datasets. This study used the genome assembly mm10 release M24 (GRCm38.p6). The publicly available datasets used in this study are listed in the methods section. Supporting data can be found in the supplementary information.

## Authors contributions

S.C. and A.P. designed the concept for this work. A.P. obtained funding and supervised the work unless otherwise stated. S.C. analysed *Srrm2* expression in *in-vivo* mouse development published datasets. S.C and T.C.C.T. grew WT and Srrm2^+/-^ mESC. S.C. performed *Mycoplasma spp.* screening. A.K. performed *Mycoplasma spp.* mapping in all datasets. S.C. performed immunofluorescence experiments. S.C. isolated RNA for RNA bulk experiments. T.C.C.T. performed qRT-PCR, western blot and alkaline phosphatase experiments for WT and Srrm2^+/-^ mESC, under the supervision of S.C. and A.P.. S.C., T.C.C.T. and A.P. interpreted the results. S.C., D.M., S.D. and A.P. designed and interpreted the flow cytometry experiments. D.M. optimized and conducted the flow cytometry experiments, under the supervision of S.D.. D.M. wrote the flow cytometry methods section of the manuscript. S.C., A.K., A.R.G. and A.P. designed the RNA-seq bulk experiments. S.C. performed RNA-seq experiments for WT and Srrm2^+/-^ mESC. S.C. and A.K. performed RNA-seq experiments for RNAi-treated mESC. P.C. performed RNA-seq mapping, quantification of transcripts and of differentially spliced events for the RNA-seq bulk datasets, under the supervision of A.R.G.. S.C. performed the computational analyses for WT and Srrm2^+/-^ mESC and for RNAi-treated mESC, under the supervision of A.R.G.. S.C. performed the qRT-PCR experiments for RNAi-treated mESC. S.C., A.R.G. and A.P. interpreted the results of the RNA-seq bulk analyses. S.C., L.Z.R. and A.P. designed the single-cell experiments, with the help of the Genomics Technology Platform of Max Delbrück Center for Molecular Medicine in the Helmholtz Association (MDC) and Berlin Institute of Health at Charité–Universitätsmedizin Berlin. S.C. and the Genomic Technology Platform performed the single-cell experiments. L.Z.R. performed the low-quality filtering, pre-processing, clustering, cell cycle assignment, and intra and inter-clusters differential expression analyses of the single-cell dataset. S.C. performed the remaining single-cell computational analyses and data visualization of all single-cell data with the help of L.Z.R.. S.C., L.Z.R., A.R.G. and A.P. interpreted the single-cell data analyses. S.C. and L.Z.R. wrote the single-cell methods section of the manuscript. S.C., P.S.B., S.G. and A.P. designed and interpreted the single-cell labeling transfer analyses. P.S.B. performed the single-cell labeling transfer analyses, under the supervision of S.G. P.S.B. and S.G. wrote the first draft of the results and methods section regarding the single-cell labeling transfer. S.C. and A.K. shared the produced datasets in the GEO repository. S.C. wrote the first draft of the manuscript, designed the figures and illustrations and produced the supplementary data. S.C., A.R.G. and A.P. wrote the final manuscript. All authors provided critical feedback and helped revise the manuscript.

## Acknowledgements

The authors thank Vedran Franke (Bioinformatics and Omics Data Science Platform, Berlin Institute for Medical Systems Biology,MDC) for advice on single-cell analyses, the System Biology Imaging facilities from the Berlin Institute for Medical Systems Biology, and the Genomics Technology Platform of Max Delbrück Center for Molecular Medicine in the Helmholtz Association (MDC) and Berlin Institute of Health at Charité–Universitätsmedizin Berlin, for their important contributions to this work, especially to Andrew Woehler, Thomas Conrad and Sarah Nathalie Vitcetz. S.C. thanks colleagues from the Pombo lab and GABBA PhD program for the productive discussions and support.

---

**Fig. S1.**
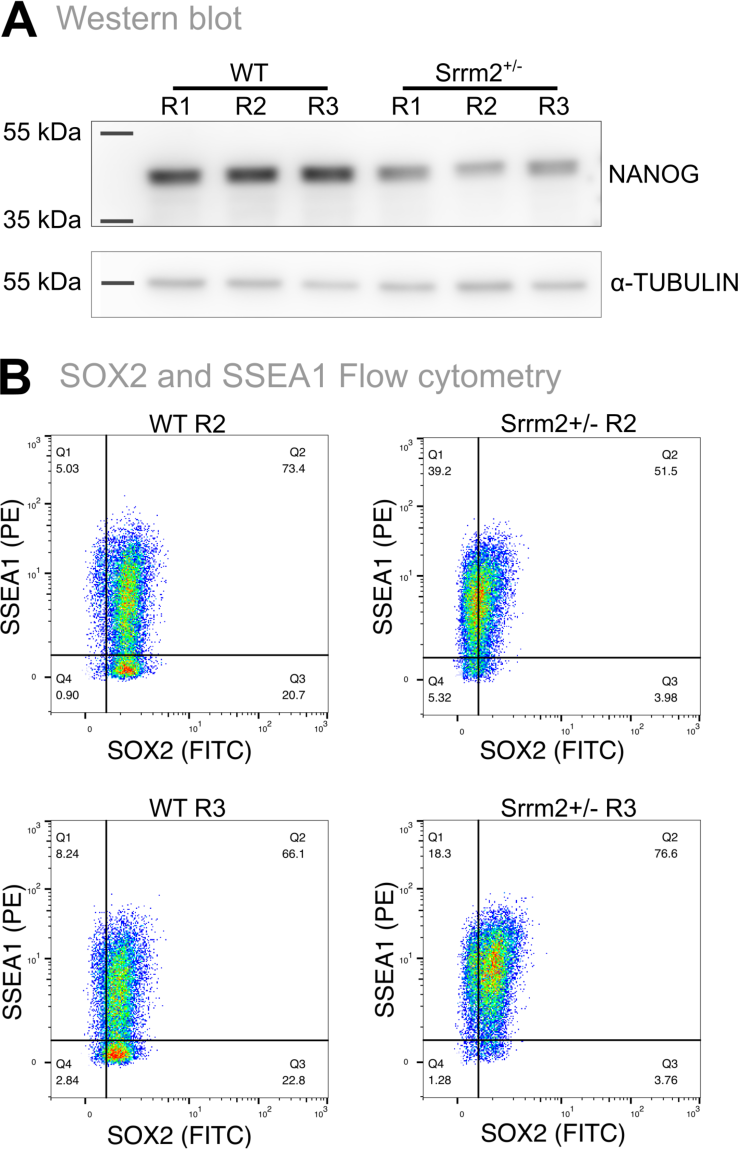
*Srrm2* expression levels influence the pluripotent state of mESC. A) Western blot analysis was performed with antibodies against NANOG and α-TUBULIN, the latter was used as a loading control. Molecular weights (kDa) are shown on the left side of each blot. Three biological replicates of WT and Srrm2^+/-^ cells are represented by different numbers on the top of the image. Data is representative of 2 independent experiments. **B)** Flow cytometry data of SSEA1 and SOX2 double labelling in WT and Srrm2^+/-^ cells. Data is representative of 2 out of 3 biological replicates, represented by different numbers.

**Fig. S2.**
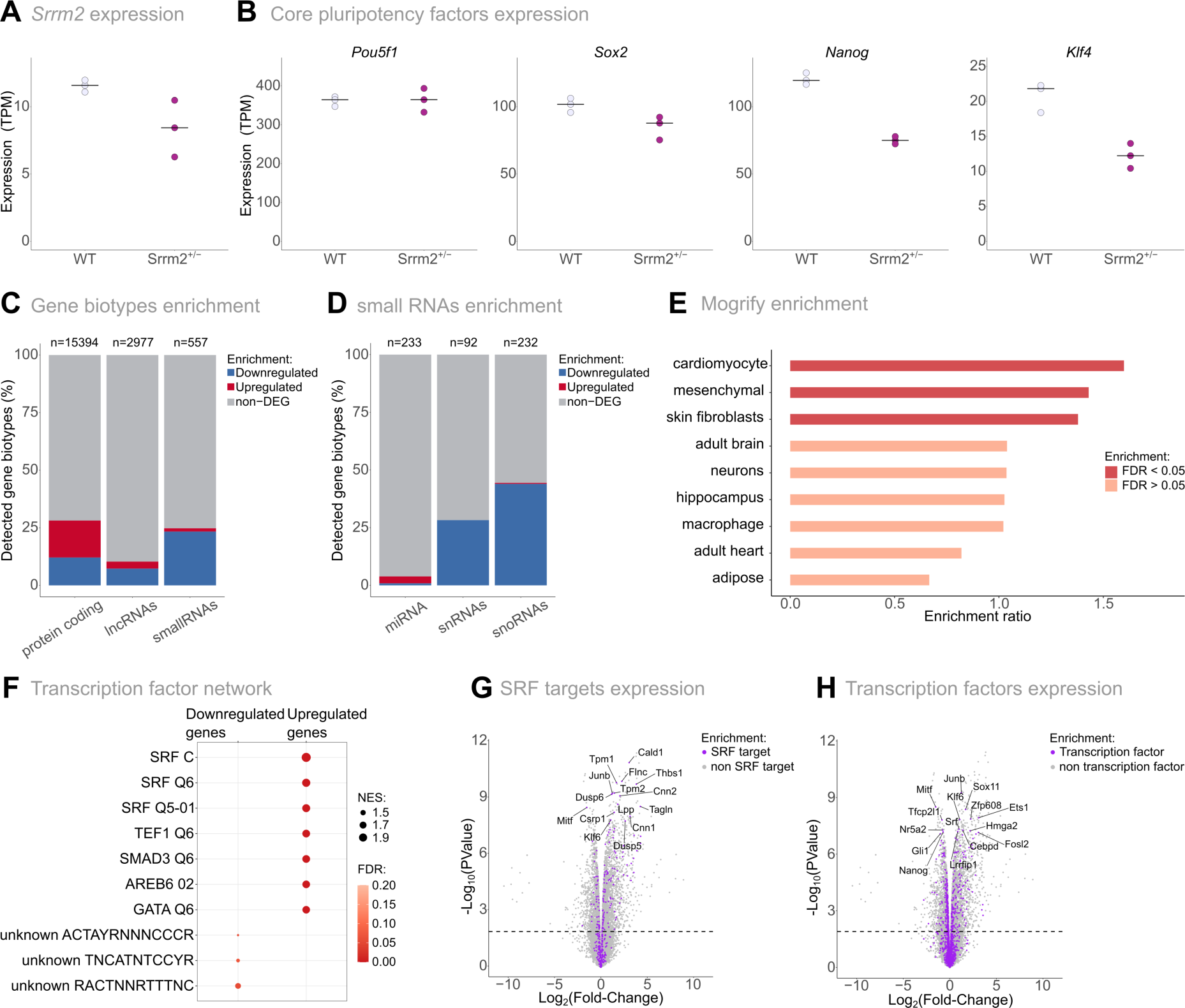
*Srrm2* heterozygous knockout mESC show extensive gene expression alterations. A) *Srrm2* expression levels in transcript per million (TPM) per replicate in WT and Srrm2^+/-^ mESC. **B)** Pluripotency transcription factors (*Pou5f1*, *Sox2, Nanog*, *Klf4*) expression levels in transcript per million (TPM) per replicate in WT and Srrm2^+/-^ mESC. **C)** Proportion of downregulated (blue), upregulated (red) and non-differential expressed genes (grey) in Srrm2^+/-^ cells per gene biotype. N, on top of each bar, represents the total number of detected genes per gene biotype: protein-coding (includes IG LV gene, TR V gene, protein-coding and TEC); lncRNAs (include Mt rRNA, Mt tRNA, antisense, misc RNA, polymorphic pseudogene, sense overlapping, processed transcript, processed pseudogene, pseudogene, rRNA, sense intronic, transcribed processed pseudogene, transcribed unprocessed pseudogene, translated processed pseudogene, unitary pseudogene, unprocessed pseudogene, lincRNA); and small RNAs (include miRNA, snRNA and snoRNA). **D)** Proportion of downregulated (blue), upregulated (red) and non-differential expressed small RNAs (grey) in Srrm2^+/-^ cells. N, on top of each bar, represents the total number of detected genes per small RNAs biotype. **E)** Over-representation analysis (ORA) for the publicly available data of transdifferentiation transcription factors imputed by Mogrify algorithm (Rackham et al., 2016; Ferrai et al., 2017) using the identified upregulated genes in Srrm2^+/-^ mESC. Different red colours represent FDR values for each enrichment term. **F)** GSEA for mouse transcription factors of Srrm2^+/-^ vs WT gene expression comparison. Red colours represent the FDR values, whereas the circle size indicates the Normalized Enrichment Score (NES) for each set of transcription factor gene targets. **G)** Volcano plot showing the expression of the SRF transcription factors targets (purple) in the transcriptional comparison of Srrm2^+/-^ and WT mESC. Gene labels represent the top 15 SRF gene targets with the lowest adjusted P-value. **H)** Volcano plot showing the expression of all *Mus musculus* transcription factors (purple) in the transcriptional comparison of Srrm2^+/-^ and WT mESC. Gene labels represent the top 15 transcription factor encoding genes with the lowest adjusted P- value.

**Fig. S3.**
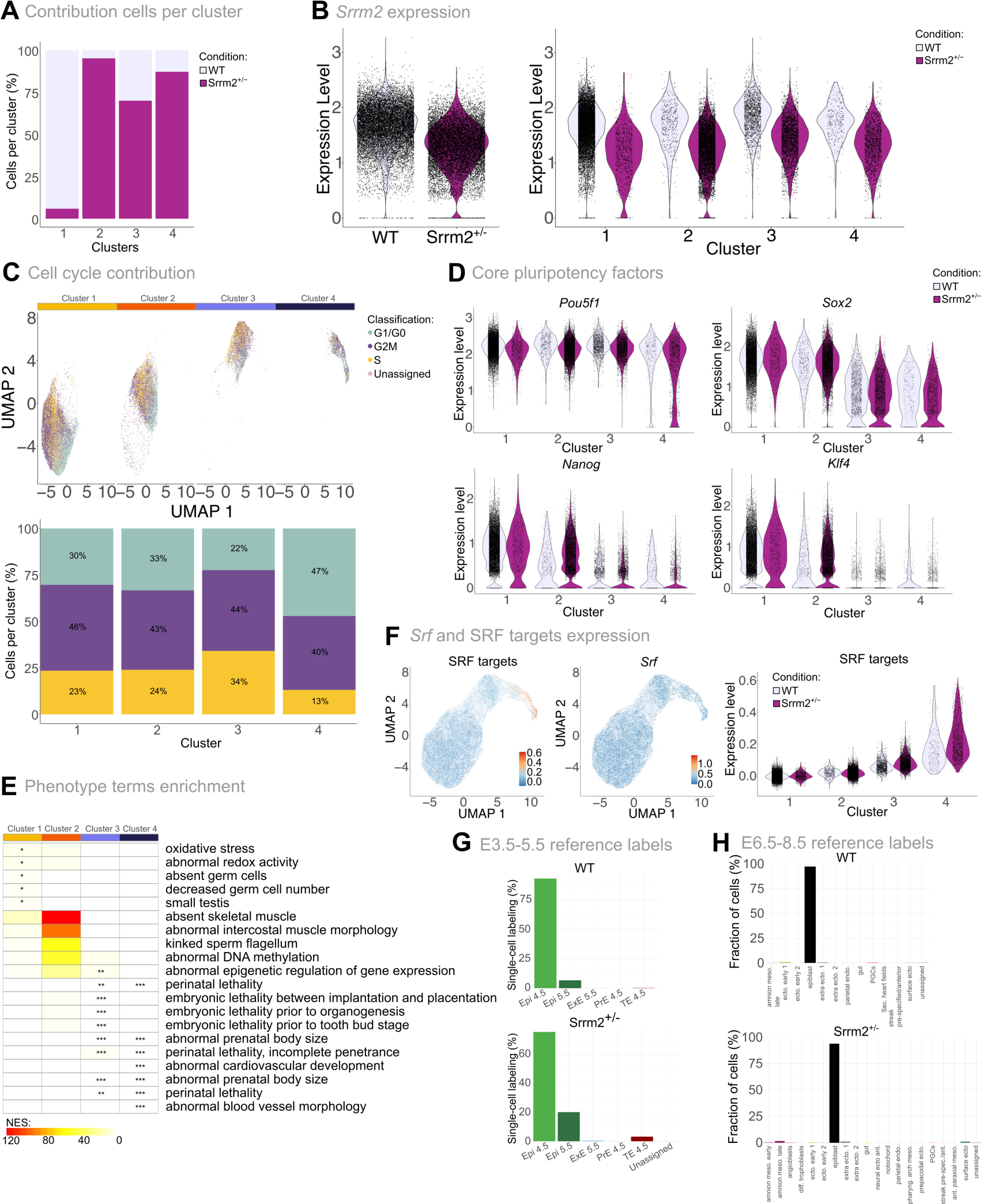
Single-cell transcriptome alterations in Srrm2 heterozygous knockout ESCs. A) Proportional contribution of WT (light pink) and Srrm2^+/-^ (dark pink) cells per cluster. **B)** Single-cell distributions of *Srrm2* expression levels per condition (left) and cluster (right). Each dot represents a single cell. Conditions are represented in different colours: WT (light pink) and Srrm2^+/-^ ESC (dark pink). **C)** Contribution of cell cycle phases. On top: UMAP projection of cell cycle phases per cell. Cells are split by cluster, identified by different colours on top. Cells are coloured by cell cycle phases. Below: Proportional contribution of cell cycle phases per cluster. **D)** Single-cell distributions of the expression levels of pluripotency transcription factors (*Pou5f1*, *Sox2*, *Nanog*, *Klf4*) per condition and cluster. Each dot represents a single cell. Conditions are represented in different colours: WT (light pink) and Srrm2^+/-^ ESC (pink). Clusters are distinguished in the y-axis. **E)** Heatmap representing phenotype terms enrichment from ORAs analyses of clusters gene markers per cluster. The top 5 significant terms per cluster are displayed. Clusters are represented by different colours on the top (from left to right, clusters 1 to 4). Normalized enrichment scores (NES) of GO terms are represented by different colours (yellow to red). Significant NES values are marked with an asterisk (Fisher exact test *: adjusted P-value < 0.05; **: adjusted P-value < 0.01; ***: adjusted P-value < 0.001). **F)** Expression of SRF gene targets or *Srf*. On the left: UMAPs representing the relative similarity between individual cells coloured by the average expression levels of SRF gene targets (left) or by *Srf* (right). Right: Single-cell distributions of SRF-target genes average single-cell expression per condition and cluster. Each dot represents a single cell. Conditions are represented in different colours: WT (light pink) and Srrm2^+/-^ ESC (pink). Clusters are distinguished in the y-axis. **G-H)** Proportion of WT (top) and Srrm2^+/-^ (down) cells assigned to predicted labels from *in vivo* scRNA-seq reference atlas of mouse embryogenesis cells from E3.5-5.5 (Nowotschin et al., 2019) (G) and E6.5-8.5 (Chan et al., 2019; Grosswendt et al., 2020, Haggerty et al., 2021) (H). G) Cells are classified into epiblast 4.5 (Epi 4.5), epiblast 5.5 (Epi 5.5), extra- embryonic ectoderm (ExE 5.5), primitive endoderm 4.5 (PrE 4.5), trophoectoderm 4.5 (TE 4.5). H) Cells are classified into amnion mesoderm early, amnion mesoderm late, angioblasts, differentiated trophoblasts, ectoderm early 1, ectoderm early 2, epiblast, extraembryonic ectoderm 1 (ExE 1), extraembryonic ectoderm 2 (ExE 2), gut, neural ectoderm anterior, notochord, parietal endoderm, pharyngeal arch mesoderm, primordial germ cells (PGCs), preplacodal ectoderm, surface ectoderm, secondary heart field/splanchnic lateral plate, splanchnic-lateral/anterior-paraxial mesoderm, streak pre-specified/anterior, unassigned (H).

**Fig. S4.**
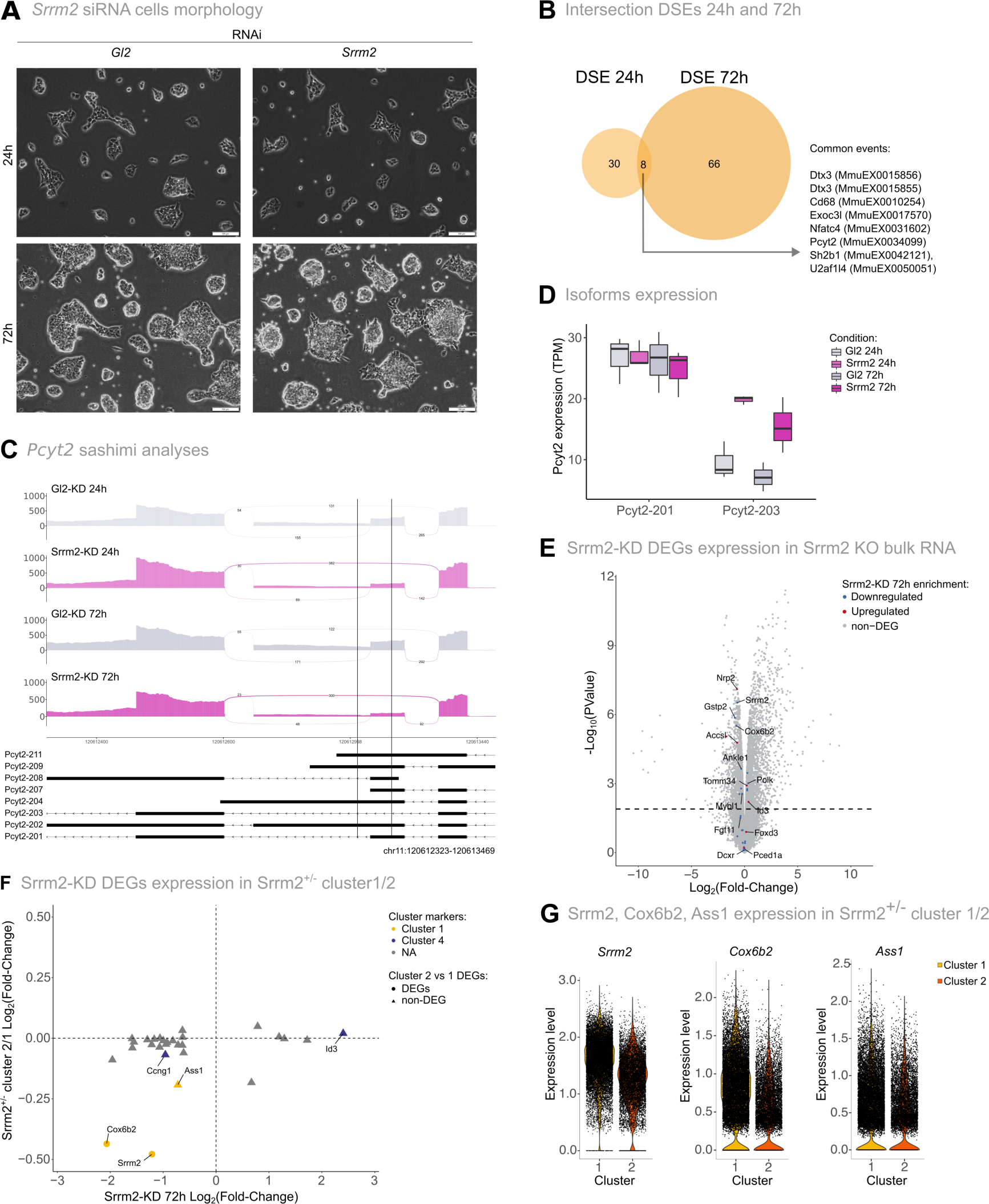
Transcriptome alterations upon *Srrm2* RNAi knockdown. A) Brightfield images from mESC cultures treated with Gl2 (control) or with *Srrm2* RNAi, at 24h or 72h. Images are representative of 4 independent experiments. Scale bar corresponds to 100 µm. **B)** Venn diagram of the differentially spliced events at 24h and 72h. The genes and the splice events of the eight common differentially spliced events between 24h and 72h are listed. **C)** Sashimi plots displaying the differential splicing events for the *Pcyt2* gene region (chr11:120612323-120613469) in control samples and Srrm2 knockdown samples at 24h and 72h. Splicing events are delimited with vertical black lines (*Pcyt2*: chr11:120613030-120613083). Numbers in the plot represent the average number of junction reads from three replicates. Isoforms that cover the represented genomic region are shown and identified as following: ENSMUST00000145356 (Pcyt2-211), ENSMUST00000142049 (Pcyt2-209), ENSMUST00000134671 (Pcyt2-208), ENSMUST00000126723 (Pcyt2-207), ENSMUST00000124468 (Pcyt2-204), ENSMUST00000106188 (Pcyt2-203), ENSMUST00000106187 (Pcyt2-202), ENSMUST00000026129 (Pcyt2-201). **D)** Expression levels (TPM) of *Pcyt2* isoforms upon Gl2 (control) and *Srrm2* knockdown at 24h and 72h. Isoforms with expression levels above 5 TPM and encoding for proteins are displayed. **E)** Volcano plot of Srrm2^+/-^ and WT transcriptional comparison with differentially expressed genes in the *Srrm2* knockdown being represented in different colours. Gene labels represent the top-10 *Srrm2* knockdown differentially expressed genes with the lowest adjusted P-value at 72h and genes mentioned in the text (*Id3*, *Nrp2*, *Fgf11*). **F)** Expression comparison (Log2 Fold change) of the differentially expressed genes upon 72h of *Srrm2* knockdown between the most stem-like single-cell clusters and *Srrm2* 72h depletion. Negative values represent downregulated genes in *Srrm2*-depleted cells and positive values represent upregulated genes in *Srrm2*-depleted cells. Gene markers of single-cell clusters are represented by different colours. Differentially expressed genes of the most stem-like single-cell clusters (cluster 2 vs cluster 1) are represented by different shapes. **G)** Single-cell distributions of the expression levels of *Srrm2*, *Cox6b2* and *Ass1* per condition and cluster. Each dot represents a single cell. Conditions are represented in different colours: WT (light pink) and Srrm2^+/-^ cells (pink). Clusters are distinguished in the y-axis.

